# PAN-CANCER ANALYSES IDENTIFY ONCOGENIC DRIVERS, EXPRESSION SIGNATURES, AND THERAPEUTIC VULNERABILITIES IN RHO GTPase PATHWAY GENES

**DOI:** 10.1101/2025.09.14.676083

**Authors:** Rubén Fernández, L. Francisco Lorenzo-Martín, Víctor Quesada, Xosé R. Bustelo

## Abstract

RHO family GTPases are key regulators of cancer-related processes such as cytoskeletal dynamics, cell migration, proliferation, and survival. Despite this, a comprehensive understanding of RHO signaling alterations across tumors is still lacking. Here, we present a pan-cancer analysis of 484 genes encoding RHO GTPases, regulators, proximal effectors, distal downstream signaling elements, and components of their proximal interactomes using data from over 10,000 tumor samples and 33 tumor types present in The Cancer Genome Atlas (TCGA). In addition, we have utilized available data from genome-wide functional dependency screens performed in more than 1,000 gene-edited cancer cell lines. This study has uncovered positively selected mutations in both well-known and previously uncharacterized RHO pathway genes. Transcriptomic profiling reveals widespread and tumor-specific differential expression patterns, some of them correlating with copy number changes. Interestingly, certain regulators exhibit consistent expression profiles across tumors opposite to those predicted from their canonical roles. Coexpression and gene set enrichment analyses highlight coordinated transcriptional programs involving some RHO GTPase pathway genes and their linkage to key cancer hallmarks, including extracellular matrix reorganization, cell motility, cell cycle progression, cell survival, and immune modulation. Functional screens further identify context-specific dependencies on several deregulated RHO GTPase pathway genes. Altogether, this study provides a comprehensive map of RHO GTPase pathway alterations in cancer and identifies new oncogenic drivers, expression-based signatures, and therapeutic vulnerabilities that could guide future mechanistic and translational research.

## INTRODUCTION

RHO GTPases can be grouped into the CDC42, RAC, RHOA, RHOD/F, RHOH, RHOU/V, RND, and the most distantly related RHOBTB subfamilies. Most of these GTPases function as molecular switches, cycling between an inactive (GDP-bound) and an active (GTP-bound) state in response to upstream stimuli. Their activity is also modulated by sequestration of the inactive forms in the cytosol. This regulatory cycle is orchestrated by three major protein classes: RHO guanine nucleotide exchange factors (GEFs), which catalyze the activation step using DBL homology (DH), DOCK or armadillo domains; RHO GTPase-activating proteins (GAPs), which promote the transition of active GTPases back to the off state using the catalytic GAP domain; and RHO GDP dissociation inhibitors (GDIs), which stabilize and sequester inactive RHO GTPases in the cytosol. Once activated, RHO GTPases engage a broad array of proximal effectors that lead to the activation of multibranched signaling cascades that influence cytoskeletal dynamics, migration, polarity, proliferation, survival, and cell type-specific functions such as lymphocyte development, T cell activation, neutrophil responses, neurogenesis, or angiogenesis among many others [1–3].

Given their wide-ranging impact on cellular physiology, most RHO GTPases have long been considered protumorigenic. This view is supported by experimental evidence demonstrating that gain-of-function mutations in some RHO GTPases, RHO GEFs or downstream effectors can drive or enhance tumorigenesis. Similarly, loss-of-function of RHO GAPs promote the same effects [1, 3–5]. Consistent with this idea, mutations in genes encoding RHO GTPase pathway elements have been found in specific cancer types such as diffuse gastric lymphoma, bladder cancer, Burkitt lymphoma, and diffuse large B-cell lymphoma (*RHOA*) [6–11]; head and neck squamous cell carcinoma (*RAC1*) [12, 13]; melanoma (*RAC1*, *PREX2*) [14–18]; and peripheral T cell lymphoma (*RHOA*, *VAV1*) [19–26] (for reviews, see [1, 3–5]). However, recent advances in cancer genomics and functional studies have revealed a more complex and context-dependent landscape for these proteins in cancer. Thus, both gain- and loss-of-function mutations in specific RHO GTPase pathway genes have been detected in human tumors, with some RHO proteins and their signaling components unexpectedly acting as tumor suppressors in certain contexts (e.g., the RHO GEFs VAV1, and TIAM1) [1, 3–5, 27–32]. Furthermore, genetic and signaling experiments have shown that some RHO signaling elements can influence tumorigenesis through both canonical (GTPase-dependent) and non-canonical (GTPase-independent) mechanisms, adding additional layers of complexity to their biological roles [1, 27]. Notably, mutations in RHO pathway genes are typically found at low prevalence in cancer genomes [1, 3–5], suggesting that altered expression levels or aberrant activation by upstream oncogenic signals may constitute their predominant mode of contribution to tumorigenesis. Despite these insights, the prevalence, diversity, and functional consequences of RHO pathway alterations across the landscape of human cancers remain poorly defined. To date, most investigations have focused on individual genes chosen based on historical precedence or laboratory interest, rather than by systematic approaches. As a result, a comprehensive, unbiased view of the mutational and transcriptional landscape of RHO GTPases, their regulators, and their downstream effectors across cancer types is still lacking.

To address this knowledge gap, we have conducted here a multidimensional analysis of 484 RHO pathway-related genes by integrating data on somatic mutations, gene expression, and copy number variation using the information available from 33 cancer types and >10,000 cancer patients that is publicly available at the TCGA [33]. In addition, we have used data from functional dependencies screens across >1,000 CRISPR–CAS9-edited cancer cell lines [34, 35]. Our findings revealed positively selected mutations in specific RHO pathway genes, a highly variegated gene expression pattern for most of them, and the identification of RHO pathway genes that are important to maintain proliferation rates in a widespread or cancer type-specific manner. Together, our study provides a foundational resource for the cancer and RHO GTPase research communities. It also reveals potential targets for further functional studies or therapeutic intervention.

## RESULTS AND DISCUSSION

### Elaboration of the list of RHO pathway-related genes

To systematically interrogate the role of RHO GTPases in cancer, we first compiled a comprehensive list of genes that have been directly or indirectly implicated in RHO GTPase-regulated signaling pathways based on biochemical, signaling, cellular, or large-scale proteomics analyses. Those included all the GTPases; direct regulators; direct effectors and distal signaling elements according to extensive literature and database searches; and proteins that form part of the large-scale RHO interactome according to proximity-dependent biotinylation (BioID)-based proteomics analyses [36]. Using these highly inclusive criteria, we ended up with a total of 484 genes encoding the following proteins (**Supplementary Table 1**): **(i)** the 23 known members of the RHO subfamily; **(ii)** the 85 proteins containing DH (2 of them also harboring RHO GAP domains), DOCK or armadillo domains that are usually involved in promoting RHO GDP/GTP exchange (2 of them also have RHO GAP domains); **(iii)** the 68 proteins that contain RHO GAP domains that are usually involved in the RHO inactivation step; **(iv)** the 3 RHO GDIs; **(v)** the 64 downstream proximal or distal effectors containing kinase domains; and **(vi)** 243 downstream proximal or distal signaling elements lacking kinase domains (e.g., regulatory factors associated with upstream receptor signaling, F-actin remodeling, cell migration, or gene transcription). This group of 484 genes will be referred to hereafter as «RHO GTPase pathway genes».

### Somatic mutational landscape of RHO pathway genes across human cancers

We used the whole-exome sequencing data from over 10,000 tumor samples associated with 33 cancer types that were contained in the TCGA database to identify the mutational landscape of RHO GTPase pathway genes. This analysis revealed that 152 (31.2%) of RHO GTPase pathway genes were mutated in more than ten cancer types, 208 (42.7%) in at least two independent cancer types, and 114 (23.4%) in specific tumor types (**Supplementary Fig. 1A**). Most of these alterations were detected at either low (<3%, 157 genes) or intermediate (3–10%, 275 genes) prevalence, although a subset of them (42 genes) exhibited mutation rates at frequencies higher than 10% (**Supplementary Fig. 1A,B**). However, since the overall mutation frequency in the foregoing analyses closely reflected both the mutational burden found in each interrogated tumor type and the size of the locus involved (**Supplementary Fig. 1B,C**), we further refined these analyses using a random sampling approach to identify the cancer types that displayed higher than expected mutation rates in RHO GTPase pathway genes when tabulated against the overall mutation burden found in the interrogated tumors. Using this approach, we found that mutations in RHO GTPase pathways genes were specifically enriched (*P* < 0.001) in 6 (18%) TCGA cancer types (BLCA, BRCA, CESC, KIRP, OV, and UCEC) and, to a lesser extent (*P* < 0.01) in 8 (24%) TCGA cancer types (GBM, HNSC, LAML, LICH, MESO, SARC, THYM, and UCS) (**Supplementary Fig. 1D**; see also **Supplementary Table 2** for the abbreviations used for the cancer type compiled in this study). In terms of prevalence, the most relevant cancer type was BRCA (*P* < 0.001) followed by KIRP, THYM, and UCEC (*P* < 0.05) (**Supplementary Fig. 1E**). These findings indicate that 14 (42%) out of the 33 TCGA cancer subclasses harbor statistically significant mutations in at least one RHO GTPase pathway gene (**Supplementary Fig. 1D,E**).

We next applied the dNdScv algorithm [37], which offers a positive selection detection method based on the normalized ratio of non-synonymous to synonymous mutations (dN/dS), to more adequately identify RHO pathway gene mutations with potential oncogenic driver roles (due to positive or negative selection patterns in specific cancer types). This analysis identified 10 RHO pathway genes (2.1% of total genes interrogated) with evidence of positive selection (qglobal < 0.01) in specific cancer types (**Fig. 1A-C**). 5 of these genes, *ACTB* (in BLCA), *RAC1* (in HNSC and SKCM), *RHOA* (in BLCA and STAD), *RHOB* (in BLCA), and *PIK3CA* (in BLCA, BRCA, CESC, COAD, ESCA, GBM, HNSC, LGG, READ, STAD, UCEC and UCS) harbored positively selected missense mutations (qmis_cv < 0.01, qtrunc_cv > 0.01) (**Fig. 1A**). Apart from *ACTB*, in which mutations have been found in Baraitser-Winter syndrome and coloboma but not in cancer [38, 39], the rest of these genes have already been detected as mutated in cancer patients in previous studies [1]. They all have been linked to protumorigenic functions, except for *RHOB* in which both tumor suppressor and protumorigenic roles have been assigned [1]. In addition, 4 out of the 10 genes identified in these analyses contained positively enriched truncating mutations (qmis_cv > 0.01, qtrunc_cv < 0.01): *ARHGAP35* (in UCEC), *MAP3K1* (in BRCA and UCEC), *SPTAN1* (in BLCA), and *SOX9* (in COAD and READ) (**Fig. 1B**). *PIK3R1* harbored both positively selected missense (in GBM and UCEC) and truncating mutations (in UCEC) (**Fig. 1A,B**), a feature that has been previously associated with bona-fide tumor suppressor genes (e.g., *TP53*). All these genes have been associated with tumor suppressor activities (*ARHGAP35*, *MAP3K1*, *PIK3R1*) [1, 40–42] or with cancer type-specific protumorigenic and tumor suppression functions (*SOX9*) [43]. Interestingly, the cancer types with positively selected mutations in *RAC1* (HNSC, SKCM) were not overlapping with those having positively selected mutations in either *RHOA* (BLCA, STAD) or *RHOB* (BLCA). However, *RHOA* and *RHOB* genes were found both mutated in BLCA (**Fig. 1A**).

**FIGURE 1.**
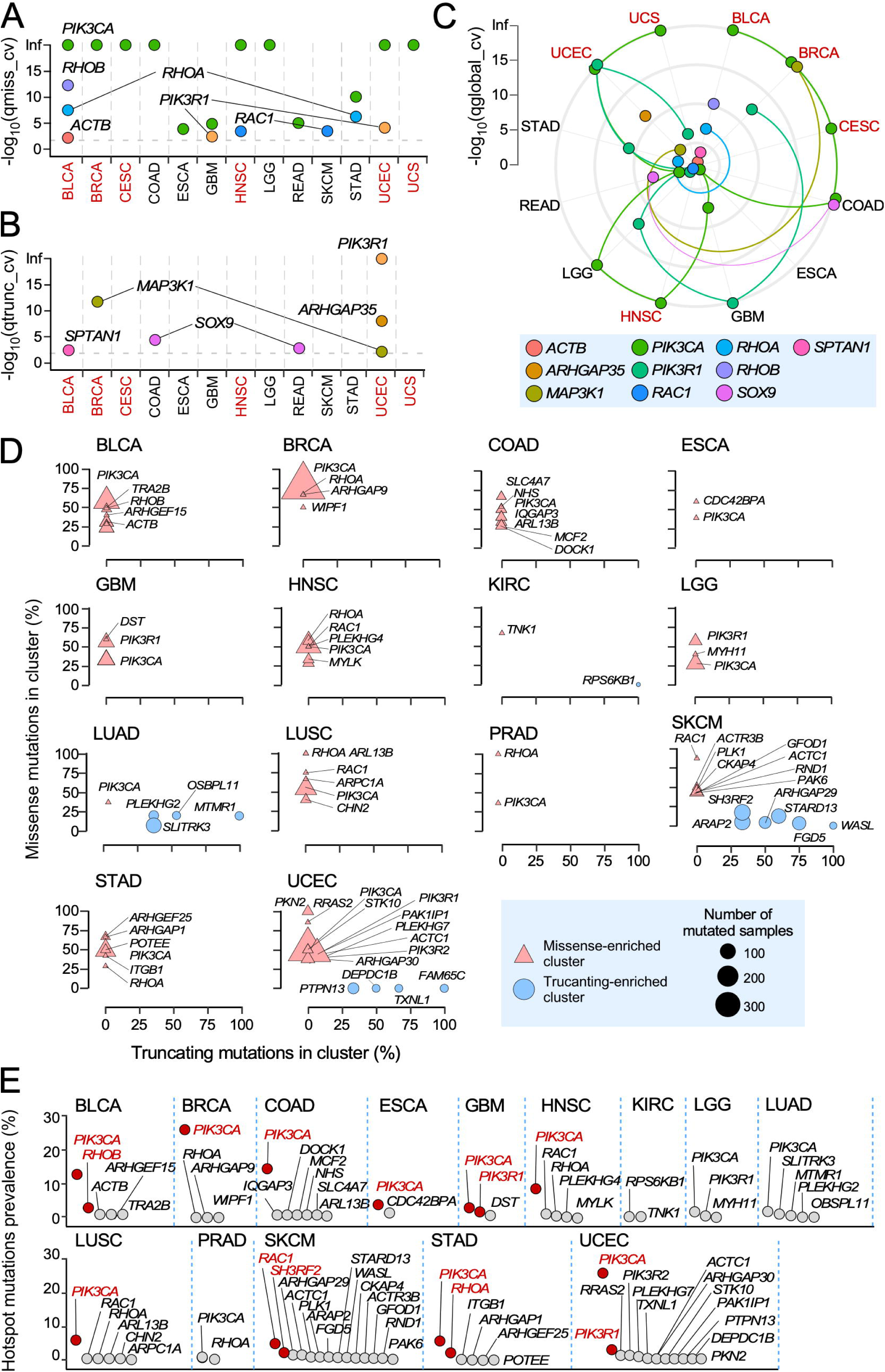
Positively selected and hotspot mutations of *RHO* pathway genes across TCGA cohorts. (**A** and **B)** Genes harboring missense **(A)** and truncating **(B)** mutations significantly associated with positive selection signals across TCGA tumors. We indicate in red tissues with enriched mutations in RHO pathway genes according to the data presented in **Supplementary Fig. 1D and E**. **(C)** Distribution of indicated positively selected mutations (bottom) across TCGA cancer subtypes according to results obtained using the dNdScv method. Each color is associated with one *RHO* pathway gene harboring somatic mutations under significant (qglobal_cv value < 0.01) positive selection. Colored lines connect the - log_10_(qglobal_cv) values associated with each *RHO* pathway gene through the different TCGA tumors. We indicate in red tissues with enriched mutations in RHO pathway genes according to the data presented in **Supplementary Fig. 1D and E**. **(D)** RHO pathway genes with hotspot mutations. Genes are stratified according to the proportion of missense (Y axis) and truncating (X axis) mutations present at the identified mutation clusters. Genes with more than 30% of their missense (triangles) or truncating (circles) mutations present at those sites are indicated by their name symbol. See inset for further information about the type and size of points used in the representation. **(E)** Prevalence of identified hotspot mutations in *RHO* pathway genes. Red dots represent genes with a hotspot mutational prevalence higher than 3% in cancer patients.

In terms of total number of positively mutated genes, BLCA (*ACTB*, *PIK3CA*, *RHOA*, *RHOB*, *SPTAN1*) and UCEC (*ARHGAP35*, *MAP3K1*, *PIK3CA*, *PIK3R1*) were the most conspicuous (4 genes with positively selected mutations each), followed by BRCA (*MAP3K1*, *PIK3CA*), COAD (*PIK3CA*, *SOX9*), GBM (*PIK3CA*, *PIK3R1*), HNSC (*PIK3CA*, *RAC1*), READ (*PIK3CA*, *SOX9*), and STAD (*PIK3CA*, *RHOA*) (2 genes with positively selected mutations each). CESC (*PIK3CA*), ESCA (*PIK3CA*), LGG (*PIK3CA*), SKCM (*RAC1*), and USC (*PIK3CA*) each contained only a positively selected gene (**Fig. 1A-C**).

As an alternative avenue to pinpoint mutations in RHO GTPase pathway genes with functionally relevant roles in tumorigenesis, we used the OncodriveCLUSTL algorithm [44] to identify hotspot mutations at specific locations of the interrogated genes. We found that 41 RHO GTPase pathway genes (8.4% of total number of interrogated genes) contained hotspot missense mutations in BLCA, BRCA, COAD, ESCA, GBM, HNSC, KIRC, LGG, LUAD, LUSC, PRAD, SKCM, STAD and/or UCEC (**Fig. 1D**, triangles). 15 out of those 41 genes showed truncation-enriched clusters that were found in a more limited number of tumor types (SKCM, LUAD, UCEC and, to a lesser extent KIRC) (**Fig. 1D**, circles). The RHO pathway genes with more positively enriched mutations included *PIK3CA* (in 12 cancers: BLCA, BRCA, COAD, ESCA, GBM, HNSC, LGG, LUAD, LUSC, PRAD, STAD, and UCEC), *RAC1* (in 4 cancers: BLCA, HNSC, LUSC, and SKCM), *RHOA* (in 4 cancers: BRCA, HNSC, PRAD, and STAD), and *PIK3R1* (in 3 cancers: GBM, LGG, and UCEC) (**Fig. 1D**). In this case, we found a cancer type (HNSC) in which hotspot mutations in *RAC1* and *RHOA* mutations were found (**Fig. 1D**). In terms of prevalence, the most relevant hotspot mutations were those present in *PIK3CA*, which were found in a large spectrum of cancer types (**Fig. 1E**, red spots). More cancer type-specific prevalent hotspot mutations were those for *RAC1* (SKCM, 4.8%), *RHOB* (BLCA, 2.9%), *PIK3R1* (UCEC, 2.7%; GBM, 2.5%), *RHOA* (STAD, 2.1%), and *SH3RF2* (SKCM, 2%) (**Fig. 1E**, red spots). These hotspot mutations included RAC1^P29S/L^ and RAC1^E31D^ (**Supplementary Fig. 2A**, top panel), RHOB^P75L/S/T^ (**Supplementary Fig. 2A**, middle panel), and RHOA^Y42S/C^, RHOA^Y34C^, RHOA^F39C^ and RHOA^E40K^ (**Supplementary Fig. 2A**, bottom panel). Hotspot missense (G376R, K379N/E) and truncation (I571Yfs*31, K575Rfs*6, T576Dfs*26, and X582_splice) mutations were also found in the case of *PIK3R1* (**Supplementary Fig. 2B**). Interestingly, the missense and truncation hotspot mutations of this gene were found segregated in GBM and UCEC, respectively (**Supplementary Fig. 2B**).

We observed a strong overlap among genes with positively selected mutations (**Fig. 1A-C**) and hotspot missense mutations (**Fig. 1D**). Such a correlation was not found in the case of the truncation mutations (**Fig. 1A-D**), a result consistent with the potential implication of these genetic alterations in loss-of-function rather than in gain- of-function events. As examples, see the distribution of these two types of mutations in *ARHGAP35*, *MAP3K1*, *PIK3R1*, and *SOX9* (**Supplementary Fig. 2B,C**). Exceptions for this lack of correlation were *PIK3R1* (which harbored mutations affecting splicing sites such as, for example, the X582_splice mutant) (**Supplementary Fig. 2B**) and *ARHGAP35*, in which the truncation hotspot R997* mutation was detected in UCEC cases (**Supplementary Fig. 2C**, top left). These results highlight distinct mutational mechanisms shaping oncogenic alterations in RHO pathway genes.

Interestingly, 7 out of the 14 cancer types previously identified as enriched in mutations in RHO GTPase pathway genes did not contain positively selected or hotspot mutations (KIRP, LAML, LIHC, MESO, OV, SARC, THYM) (**Fig. 1A-C** and **Supplementary Fig. 1D,E**; cancer types in black font; **Supplementary Table 3**). Conversely, there were 7 cancer types that contained positively selected or hotspot mutations that did not have a differential enrichment in mutations for RHO GTPase pathway genes (COAD, ESCA, GBM, LGG, READ, SKCM, STAD) (**Fig. 1A-C**, **Supplementary Fig. 1D,E**, and **Supplementary Table 3**). In contrast, there were 6 cancer types that shared both positively selected and hotspot mutations (BLCA, BRCA, CESC, HNSC, UCEC, and UCS) (**Supplementary Fig. 1D,E**; **Fig. 1A-C**, tumors shown in red font; **Supplementary Table 3**). In the case of cancer type overlap, we found RHO GTPase pathway genes that displayed positively enriched and hotspot mutations in the same cancer types (e.g., *RAC1* in HNSC and SKCM; *RHOA* in STAD; *RHOB* in BLCA; *PIK3CA* in BLCA, BRCA, ESCA, GBM, HNSC, LGG, STAD, and UCEC; *PIK3R1* in GBM and UCEC) (**Fig. 1B-D**). However, this overlap was not seen in all cases (e.g., *PIK3R1* in LGG; *PIK3CA* in COAD, *RHOA* in BCLA) (**Fig. 1B-D**). In contrast, we found little overlap between selected mutations and hotspot mutations likely due to the reasons stated above (**Fig. 1B-D**). Overall, the hotspot analyses identified 4-fold more RHO pathway genes with potential oncogenic driving roles than those based on the detection of positively selected mutations (**Fig. 1B-D**). Interestingly, these analyses also indicated that there were 13 TCGA cancer types that did not contain positively selected or hotspot mutations (ACC, CHOL, KICH, KIRC, LUAD, LUSC, DLBC, PAAD, PCPG, PRAD, TGCT, THCA, and UVM) (**Supplementary Table 3**).

### Transcriptional deregulation landscape of RHO pathway genes in human cancers

We next explored the degree of deregulation of RHO pathway genes at the mRNA level. To this end, we performed gene expression analyses across 20 TCGA cancer cohorts that included matched normal tissue controls. Overall, we found that 358 (74%) of the 484 interrogated RHO pathway genes were deregulated in a statistically significant manner in at least one cancer type (**Fig. 2A,B**). The extent and type of deregulation of the transcripts, however, varied substantially across the different tumor types (**Fig. 2A,B**). For example, >40% of RHO pathway genes were differentially expressed in LIHC, LUSC and GBM, while <3% were found in SARC and THYM (**Fig. 2B**). In addition, we found that upregulation events were highly predominant in LIHC, whereas in other cases downregulation events prevailed (GBM, KIH, PRAD, THYM, SARC) (**Fig. 2B**). Random sampling analyses confirmed that these alterations occurred at significantly higher rates than expected by chance in all the 20 interrogated TCGA cancer types (**Fig. 2C**), suggesting that transcriptional deregulation of RHO signaling is a recurrent and biologically relevant feature of tumorigenesis. Cancer types with differentially expressed RHO GTPase pathway genes included those that were also enriched in overall mutations, in positively selected mutations, and in hotspot mutations (**Fig. 2C**, cancer types shown in red font) as well as two TCGA cancer types (KICH, THCA) that were not identified in our previous mutational analyses (**Fig. 2C**, cancer types shown in black font). The transcripts for RHO GAPs were the most frequently deregulated (86.8%), followed by downstream kinases (81.3%), RHO GEFs (81.0%), RHO GTPases (78.3%), RHO GDIs (66.7%), and downstream elements lacking kinase activity (65.0%) (**Fig. 2D**). However, a minority of RHO GTPase pathway genes was consistently upregulated (**Fig. 2D**, red color) or downregulated (**Fig. 2D**, blue color) in all cancer types. RHO GAP-encoding genes were by far the RHO GTPase pathway genes with more consistent patterns of pan-cancer upregulation or downregulation (58.8% of interrogated RHO GAPs), followed by kinase interactors (45.3%), non-kinase interactors (40.7%), RHO GEFs (31.0%), RHO GTPases (21.8%), and RHO GDIs (0%). The functional subclasses with lower numbers of differentially expressed events across cancers were non-kinase interactors (35%) and RHO GDIs (33.3%) (**Fig. 2D**). The expression of RHO GAP-, RHO kinase interactor-, RHO GEF-, and RHO GTPase-encoding genes did not change in 13.2%, 18.7%, 19.0%, and 21.7% of cases, respectively (**Fig. 2D**).

**FIGURE 2.**
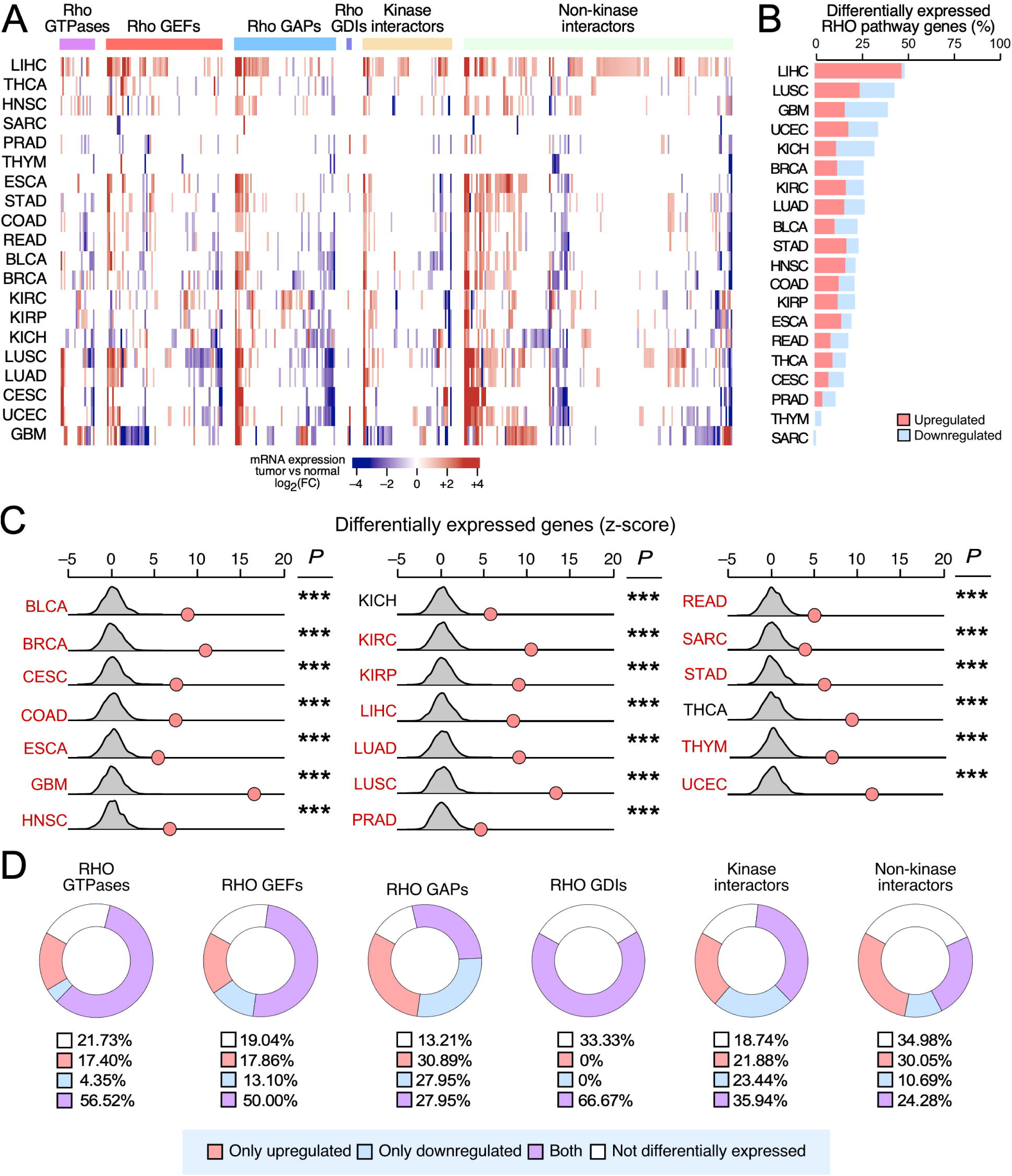
Differential expression of RHO pathway genes in pan-cancer data. **(A)** Heatmap showing the RHO pathway genes (top) that are either upregulated (red color) or downregulated (blue color) in indicated cancer types (left). Expression is presented as a gradient from most downregulated to most upregulated genes (bottom). **(B)** Stacked bar diagram representing the significantly up- (red) and downregulated (blue) RHO pathway genes across TCGA tumors. **(C)** Results from the random sampling analysis of RHO pathway genes differential expression across TCGA tumors, comparing the number of differentially expressed RHO pathway genes (red dots) with an expected background distribution of unrelated genes (grey shapes). *P* were calculated using the Poisson test: *, *P* < 0.05; **, *P* < 0.005; ***, *P* < 0.0005. **(D)** Circular stacked plots representing the percentage of RHO pathway genes in the indicated expression subclasses (blue inset at the bottom).

In terms of penetrance, we found that 44% of deregulated RHO pathway genes were differentially expressed in more than 5 TCGA cancer types and defined this subset as «pan-cancer differentially expressed RHO GTPase pathway genes» (**Fig. 3A,B**). On the other hand, some RHO GTPase pathway genes showed a more limited spectrum of differential expression across cancer types, including some whose deregulated expression was only found in one (20%) (**Fig. 3C-G**), two (14%), three (14%) or four (8%) cancer types (see examples in Fig. **Fig. 3C** to **I**). Consistently upregulated transcripts encoding RHO GTPase pathway elements across 25-50% (not underlined in the text) and >50% (underlined in the text) of the 20 interrogated TCGA cancer types included 4 RHO GTPases (*<u>RAC3</u>*, *RAC2*, *RHOD, RHOV*), 9 RHO GEFs (*<u>ECT2</u>*, *<u>PLEKHG4</u>*, *ARHGEF16, ARHGEF19*, *ARHGEF38*, *DOCK6*, *FGD6*, *PLEKHG2*, and *VAV2*), 12 RHO GAPs (*<u>ARHGAP11A</u>*, *<u>DEPC1</u>*, *<u>DEPC1B</u>*, *<u>RACGAP1</u>*, *ARHGAP11B*, *ARHGAP11B*, *ARHGAP22*, *ARHGAP33*, *ARHGAP39*, *ARHGAP4*, *ARHGAP8*, and *SH3BP1*), 4 downstream kinases (*<u>CIT</u>*, *<u>LIMK1</u>*, *<u>PLK1</u>*, and *PLK2*), and 24 non-kinase downstream elements (*<u>ANLN</u>*, *<u>DIAPH3</u>*, *<u>IQGAP3</u>*, *<u>KIF14</u>*, *<u>LMNB1</u>*, *<u>RHPN1</u>*, *<u>RTKN</u>*, *<u>SOX9</u>*, *ACTR3B*, *AMIGO2*, *ARPC1B*, *BAIP2L1*, *CCT2*, *CCT6A*, *DSG2*, *EPSTI1*, *HSPE1*, *SCRIB*, *SHMT2*, *SLC1A5*, *TFRC*, *TMPO*, *TUBA1B,* and *VANGL2*) (**Fig. 3A,B**). Consistently downregulated transcripts across >50% (underlined in the text) and 25-50% (not underlined in the text) of the analyzed cancer types included 2 RHO GTPases (*<u>RHOJ</u>*, *RHOB,* and *RHOU*), 4 RHO GEFs (*<u>ARHGEF15</u>*, *ARHGEF17*, *ARHGEF6*, and *FGD5*), 10 RHO GAPs (*<u>ARHGAP20</u>*, *<u>DLC1</u>*, *ARHGAP10*, *ARHGAP24*, *ARHGAP28*, *ARHGAP31*, *MYO9A*, *STARD8*, *STARD13*, and *SYDE2*), 2 downstream kinases (*<u>MYLK</u>* and *RPS6KA2*), and 9 non-kinase downstream elements (*<u>AKAP12</u>*, *<u>FERMT2</u>*, *<u>MYH11</u>*, *<u>WASF3</u>*, *CDC42EP2*, *FAM65B*, *FNBP1*, *SLITRK3*, and *SPTBN1*) (**Fig. 3A,B**). These results suggest that these consistent expression patterns across cancers might have functional relevance. In line with this, we found that 5 of those upregulated genes (*DEPC1B*, *IQGAP3*, *PLEKHG2*, *PLK1*, and *SOX9*) and 6 of those downregulated genes (*FGD5*, *MYLK*, *MYH11*, *RHOB*, *STARD13*, *SLITRK3*) also harbored positively selected and/or hotspot mutations according to previous analyses (see above, **Fig. 1**). However, many RHO GTPase pathway genes showed different patterns of expression (up- and downregulation) depending on the cancer type analyzed (**Figs. 2C** and **3A-H**). This is the case, for example, for 2 RHO GTPase-encoding genes (*RHOF* and *RND2*) (**Fig. 3B**), 9 RHO GEF-encoding genes (*<u>PREX2</u>*, *ARHGEF4*, *DOCK3*, *DOCK11*, *PLEKHG4B*, *PLEKHG5*, *RASGRF1*, *RASGRF2*, *VAV3*) (**Fig. 3A,B**), 2 RHO GAP-encoding genes (<u>ARHGAP6</u>, *TGAP*) (**Fig 3A,B**), 5 kinase encoding genes (*CKB*, *DGKA*, *DGKG*, *DMPK*, *PAK6*) (**Fig. 3B**), and 10 genes encoding either non-kinase downstream or interactome elements (*<u>CAV1</u>*, *FAM169A*, *FAM15C*, *IQGAP2*, *MCAM*, *MUC13*, *NCF2*, *NHS*, *PCDH7*, and *RNKN2*) (**Fig. 3A,B**). This cancer type-specific expression pattern could represent bystander events without any functional relevance or the fact that some of those genes encode proteins with dual functions depending on the cancer context. Alternatively, it might reflect the enrichment of a specific cell population that originates the cancer and that has expression levels of RHO pathway genes different from the rest of cell types that form part of that tissue from which the cancer has originated from. In general, the most consistent expression patterns were associated with RHO GTPase pathway genes that showed differential expression patterns in more than 50% of the interrogated cancer types (**Fig. 3A**).

**FIGURE 3.**
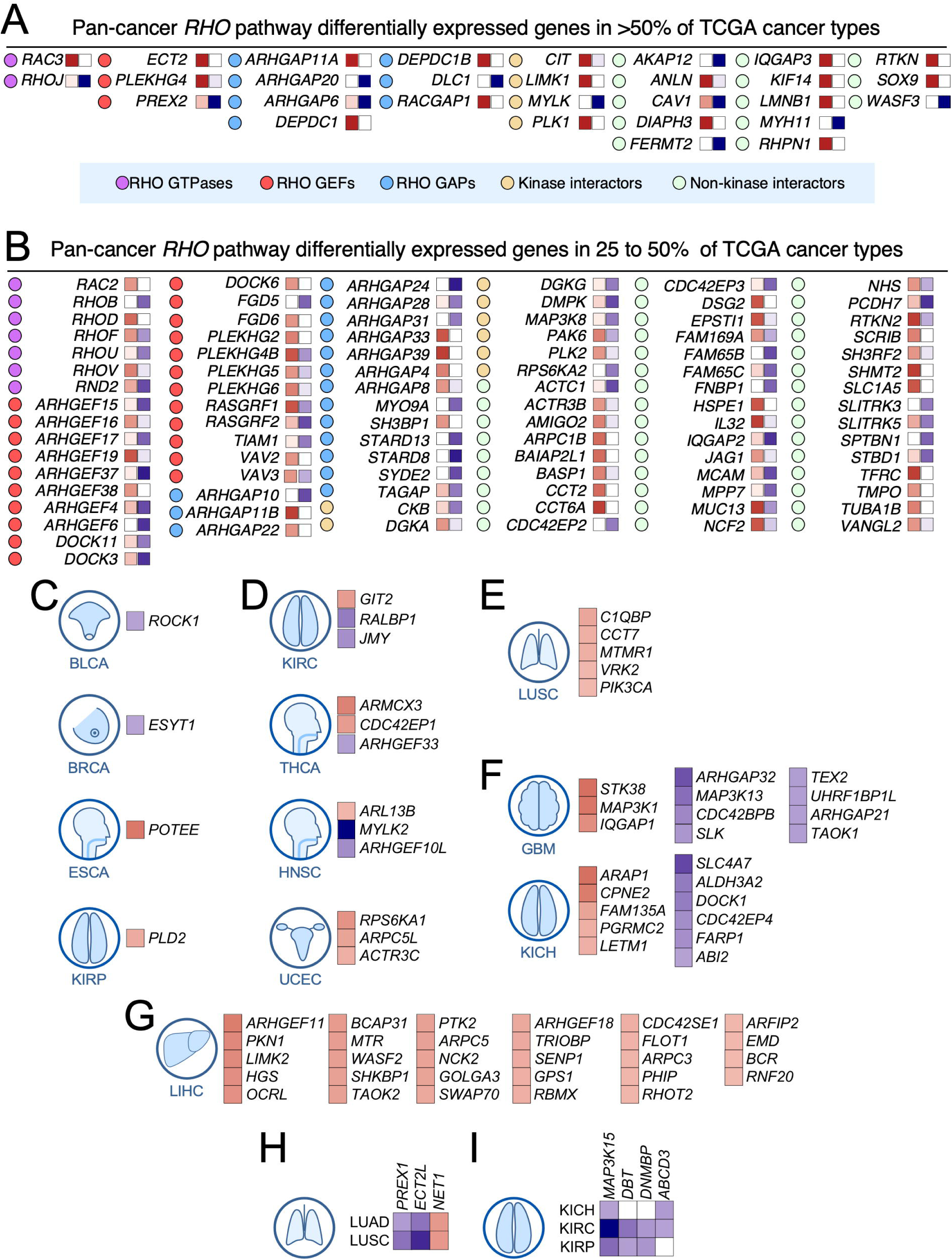
Differential expression of RHO pathway genes in pan-cancer data. **(A** and **B)** RHO pathway genes that are significantly up-(red) or downregulated (blue) in more than 50% (A) or 25% (B) of the analyzed TCGA cohorts. The brightness of squares color is directly proportional to the total number of tumors in which each gene is significantly up- or down-regulated. Tumors associated with brighter red than blue were considered pan-cancer upregulated *RHO* pathway genes, and vice versa. **(C** to **I)** Examples of RHO pathway genes with more expression patterns limited to one (C to G), two (H), and three (I) tumor types. For expression color codes, see panels A and B.

These analyses also indicated that many RHO pathway elements did not follow the expected transcriptional deregulation behavior according to their canonical functions (**Figs. 2C** and **3**, **Supplementary Fig. 3**). For example, RHO GEFs, which act as positive regulators of RHO GTPases, were expected to be upregulated in most tumor types. However, this expected expression pattern only was clearly observed in some RHO GEFs (*ARHGEF11*, *ARHGEF18*, *ARHGEF19*, *DOCK6*, *ECT2*, *FGD6*, *NET1*, *PLEKHG2*, and *VAV2*) (**Fig. 3A,B,G,H**; **Supplementary Fig. 3**). Consistent protumorigenic functions for some of those RHO GEFs (*ARHGEF11*, *ARHGEF18*, *ARHGEF19*, *ECT2*, *PLEKHG2*, and *VAV2*) have been described in some tumor models in previous studies [1, 45–51]. Likewise, RHO GAPs, which act as negative regulators of RHO GTPases, were supposed to be downregulated in most tumor types. However, this was only the case of some RHO GAPs such as *ARHGAP20*, *ARHGAP21*, *ARHGAP24*, *ARHGAP31*, *ARHGAP32, DLC1*, *MYOA9*, *STARD8*, *STARD13*, and *SYDE1* (**Fig. 3A,B**,**F**; **Supplementary Fig. 3**). Tumor suppressor-like functions for some of those RHO GAPs have been described (ARHGAP20, ARHGAP24, DLC1, MYOA9, STARD8, and STARD13) [1, 52–59]. However, in contrast to the expected pattern of expression, we found another subset of RHO GEFs (*ARHGEF4*, *ARHGEF6*, *ARHGEF15*, *ARHGEF17*, *ARHGEF37*, *FGD5*, *PREX2, TIAM1, RASGRF2*) and of RHO GAPs (*ARHGAP4*, *ARHGAP8*, *ARHGAP11A*, *ARHGAP11B*, *ARHGAP22*, *ARHGAP33, ARHGAP39*, *DEPC1*, *DEPC1B*, *RACGAP1, SH3BP1*) that were unexpectedly downregulated and upregulated, respectively (**Figs. 2D** and **3A,B,D,F-H**; **Supplementary Fig. 3**). This suggests that a significant fraction of RHO GEFs and RHO GAPs play opposite functions to the expected canonical ones. In addition, these data suggest that the mutations found in some of those RHO GEFs (*ARHGEF15*, *FGD5*) and RHO GAPs (*DEPC1B*) in previous analyses (**Fig. 1**) might be associated with loss-of-function events. This unexpected pattern of expression can be due to several causes. Firstly, that the encoded proteins can play catalysis-independent functions as previously shown for specific RHO GTPase pathway elements. Alternatively, they can participate in the regulation of key biological processes for cancer cells. This is the case, for example, of the essential involvement of RACGAP1 in cytokinesis and other pro-tumorigenic pathways [60, 61]. Consistent with this idea, we have found that 3 of the upregulated RHO GAP-encoding genes identified in our *in silico* analyses (*ARHGAP11A*, *ARHGAP11B*, *DEPDC1*; **Fig. 3A,B**) do perform RACGAP1-like functions in cytokinesis (unpublished data). Further agreeing with this hypothesis, previous studies also have demonstrated protumorigenic roles for both ARHGAP11A and RACGAP1 in basal-like breast cancers [62]. Alternatively, the noncanonical deregulation of the expression levels of RHO GTPase pathway genes might indicate that such changes represent just a bystander, functionally irrelevant event (due to transcriptional programs or expansion of the cell type that originates the final tumor mass). Cases of opposite spectra of mRNA expression have been also found for the rest of functional subclasses of RHO GTPase pathway genes studied in this work (**Figs. 2D** and **3A,B**; **Supplementary Fig. 3**). Future wet-lab analyses will have to be done to decipher these issues.

To further explore the biological basis of the differential expression patterns of RHO GTPase pathway genes, we investigated whether such an expression was associated with the catalytic specificity towards their GTPase substrates. To this end, we used the proposed list of substrates (CDC42, RAC1 and RHOA) for these RHO GTPase regulators that was previously generated based on information gathered from fluorescence resonance energy transfer-based activity assays [63]. Among the 12 identified pan-cancer downregulated RHO GAPs, we found that 50% of them were specific for RHOA, while only one (SYDE2) was considered RAC1-specific. In contrast, 36% of the eleven upregulated RHO GAPs were specific for RAC1 while only two (*ARHGAP8* and *ARHGAP11B*) were proposed to be RHOA-specific (**Supplementary Fig. 4**). Given that there is a similar number of RHOA- and RAC1-specific RHO GAPs encoded in the human genome [63], these results suggest that many cancer types downregulate the expression in favor of RHOA activation and RAC1 inactivation with the downregulation of RHOA GAPs and the upregulation of RAC1 GAPs. In the case of RHO GEFs, we found that the upregulated subset primarily contained RHOA-(50%) and CDC42-specific (18.2%) GEFs (**Supplementary Fig. 4**). In contrast, the downregulated subset contained a 50% of RAC1-specific GEFs (**Supplementary Fig. 4**). However, it is worth noting that in this case these data might not reflect changes associated with a functional trend, since the human genome approximately contains a 7:1 ratio between RHOA and RAC1-specific GEFs [63]. These results suggest that some cancer types favor the activation of RHOA via downmodulation of the expression of RHOA GAP-encoding genes rather than through the deregulation of the expression of RHO GEFs.

### Impact of copy number variations on the expression of RHO pathway genes

Somatic copy number variations (SCNVs) can also contribute to changes in gene expression. To explore this possibility, we used a new algorithm developed by us (CiberAMP) that can establish direct correlations between gene copy number changes and significant shifts in mRNA expression levels [64]. As a first step, we found using CiberAMP that 87% of the 484 RHO GTPase pathway genes undergo SCNVs events in at least one of the 33 interrogated TCGA cancer types. When focusing on SCNVs with high prevalence (affecting >10% of patients), this proportion decreases to 74% (**Fig. 4A**). Shallow amplifications were the most frequent SCNVs across TCGA tumors, followed by deep amplifications, and shallow and deep deletions, respectively (**Fig. 4A).** To distinguish driver events from passenger alterations co-occurring with known oncogenes, we used CiberAMP to identify the SCNVs in RHO pathway genes that mapped in close vicinity with well-established oncogenes listed in the COSMIC Cancer Gene Census (CGC). We found that 97% of SCNVs associated with RHO GTPase pathway genes do significantly co-occur with amplifications or deletions of CGC oncogenes (**Fig. 4A**, light colored circles), indicating that most SCNVs detected for RHO GTPase pathway genes are probably mere passenger events in these tumors. In line with this, we found using CiberAMP that 70% of all the RHO GTPase undergoing deep (**Fig. 4B**) or shallow (**Fig. 4C**). SCNVs did not change their expression according to the gene loss or gene amplification event detected. Furthermore, we could only identify 10 pan-regulated RHO pathway genes in which the gene losses (*MYH11*) or gene amplifications (*ANLN*, *ARHGAP39*, *DEPDC1*, *ECT2*, *IQGAP3*, *KIF14*, *PLEKHG4B*, *RHPN1*, *TFRC*) did consistently reinforce (>80% of the cohorts in which SCNVs were detected) the transcriptional changes found in analyses performed using cancer samples vs healthy controls (**Fig. 4D**). Two of those genes also undergo hotspot mutations (*MYH11*, *IQGAP3*) according to our prior analyses (**Fig. 1D**). Although small in number, changes in the expression of these genes must be relevant, given that many of them have been linked to either tumor suppressor (*MYH11*) or tumor promoting (rest of them) functions (see also below, **Fig. 6B**). Taken together, these data indicate that only a minority of RHO GTPase pathway genes seem to change their expression because of SCNVs.

**FIGURE 4.**
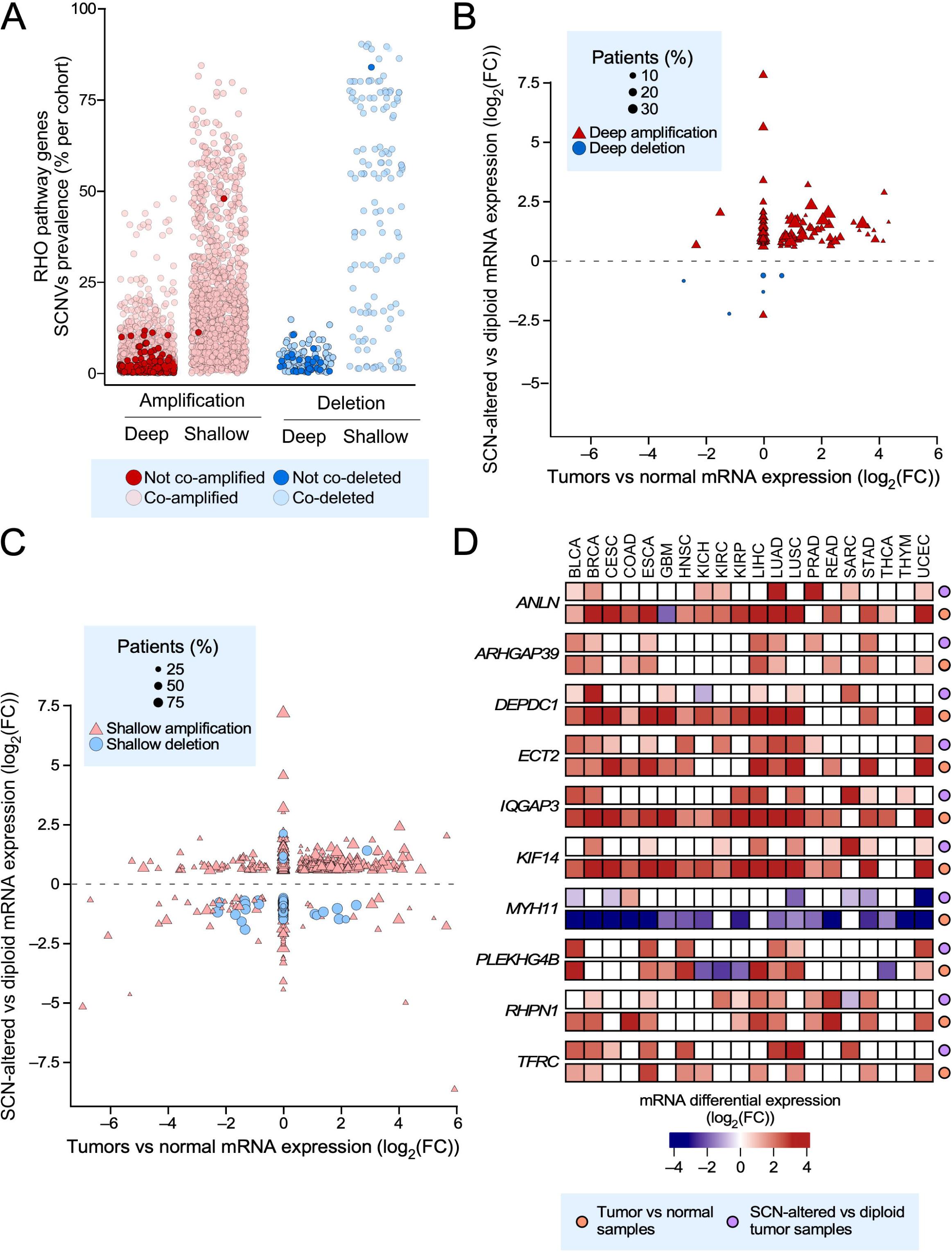
Somatic copy number variations associated with transcriptional deregulation of RHO pathway genes in TCGA tumors. **(A)** RHO pathway SCNVs found in pan-cancer data. Dot colors represent copy number gains (red) and losses (blue). Dark colors represent copy number variations of RHO pathway genes that are not significantly coamplified or codeleted with any known cancer driver present in the COSMIC database. **(B** and **C)** Transcriptional status analysis of SCN-associated RHO pathway differentially expressed genes associated with highly deep (B) or shallow (C) SCNVs (> 10% of the patients). Y axis represents the log_2_(FC) value of RHO pathway genes calculated during the comparison of their mRNA levels between SCN-altered and diploid tumor tissues. X axis represents the log_2_(FC) value of RHO pathway genes calculated by comparing the mRNA level variations found between tumor and normal samples. Dot size is proportional to the prevalence of copy number variations. **(D)** Heatmap representation of the log_2_(FC) values associated with the differential expression of 10 pan-cancer deregulated RHO pathway genes calculated during the comparison of their mRNA levels between SCN-altered and diploid tumor tissues (purple dots) and tumor vs normal samples (orange dots) in TCGA tumors.

### Coordinated pan-cancer deregulation of RHO-specific GEFs and GAPs

Given our observation that many RHO GEFs and GAPs are consistently deregulated across multiple tumor types (**Fig. 3A,B**), we next explored whether such expression patterns could represent their participation in large scale coordinated transcriptional programs in specific cancer types. To assess this issue, we performed coexpression analyses between all possible pairs of RHO GEFs, RHO GAPs, and RHO GEFs–RHO GAPs across the 33 TCGA cohorts to find coregulatory events. These analyses revealed that 98% of all possible RHO GEF–RHO GAP gene pairs show significant coexpression in >25% of interrogated tumor types (**Fig. 5A,B**). The most relevant coregulated GAP–GEF pairs (>50% of cancer types) included *ARHGAP11A–ECT2*; *MYO9A* with *ARHGEF17* and *PLEKHG5*; *ARHGAP6*–*ARHGEF17*; *SH3BP1*–*TIAM1*; and *STARD13*–*PLEKHG5* (**Fig. 5A**, purple color dots) while the most relevant anti-regulated GAP–GEF pairs included those of *ECT2* with *ARHGAP10* and *STARD13*; and *RACGAP1–FGD5* (**Fig. 5A**, green color dots). We also found cases of strong coregulation among RHO GAP pairs in pan-cancer data, including *SH3BP1* with *RACGAP1* and *ARHGAP6*; *STARD13* with *ARHGAP6, ARHGAP11A,* and *MYO9A*; and *ARHGAP10–MYO9A* (**Fig. 5B**, purple color dots) as well as cases of strong antiregulation (*STARD13–ARHGAP11A*; *ARHGAP10–MYOA9*; and *ARHGAP8–TGAP*) (**Fig. 5B**, green purple dots). In contrast, the pattern of strong coregulation (*ARHGEF19–PLEHHG5*) and anti-regulation (*ECT2–VAV3*) among RHO GEFs was much more limited (**Fig. 5C**, purple and green dots, respectively). Although limited to reduced number of tumors, the *ARHGEF17–PLEKHG5* pair also showed a high statistical significance in 40% of the cancer types interrogated (**Fig. 5C**). Additional pairs with lower percentages of distribution in cancer types (less than 50%) were found for all those categories (**Fig. 5A-C**).

**FIGURE 5.**
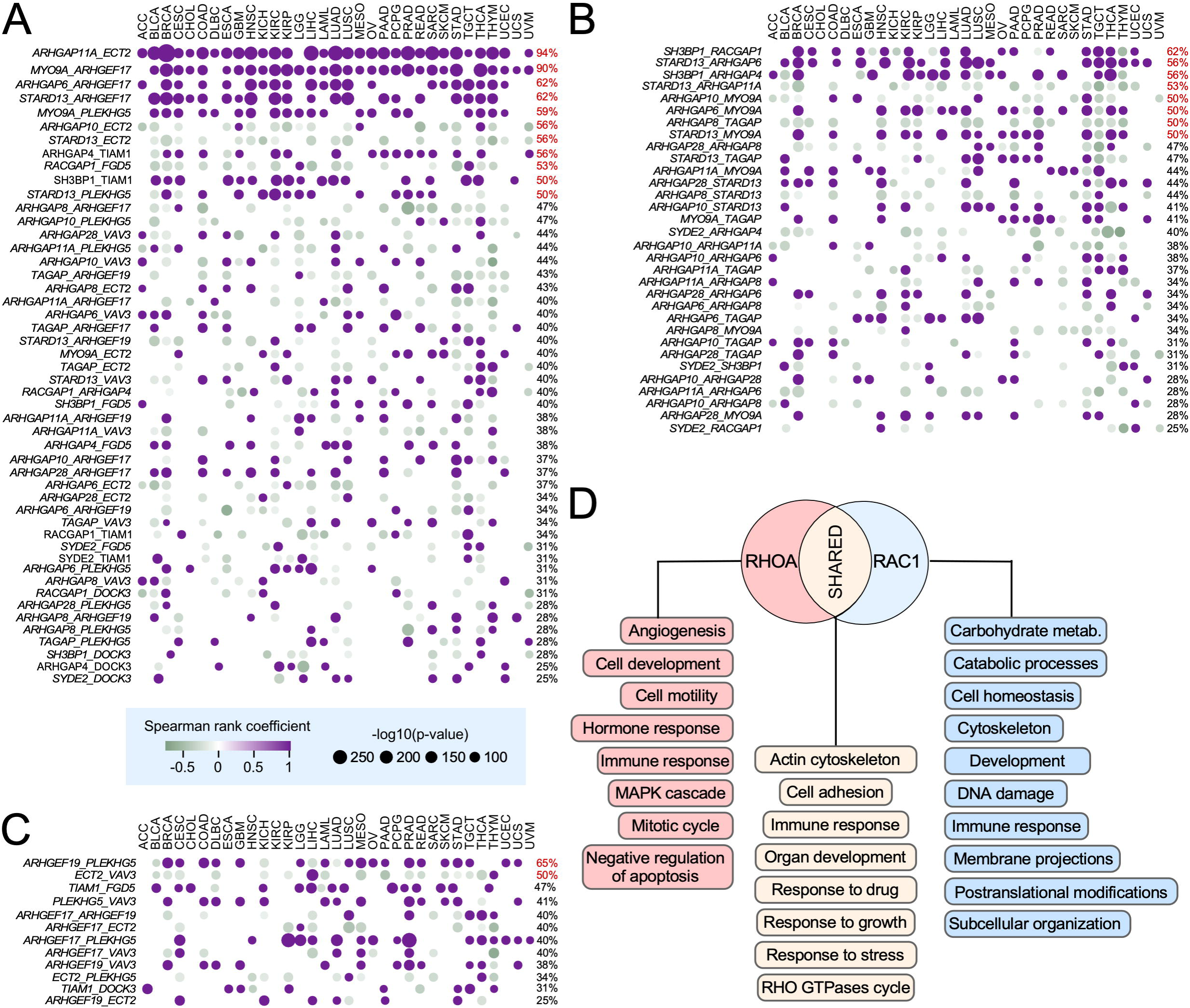
Ontological analysis of pan-cancer deregulated RHO GEFs and GAPs. **(A** to **C)** Coexpression analysis of indicated pairs of RHO GAPs–RHO GEFs (A, left), RHO GAPs (B, left), and RHO GEFs (C, left) in indicated tissues (A, B and C, top). Dot size correlates with each pair associated -log_10_(*P*), and their color is correlated Spearman correlation coefficient (positive correlations in purple, and negative correlations in green). The percentage of TCGA tumors in which the expression of the indicated pair of genes is significantly coregulated is indicated on the right. In red color, those with coexpression detected in more than 50% of cancer types. **(D)** Top global GO terms associated with RHOA- and RAC1-specific pan-cancer deregulated GEFs and GAPs across TCGA tumor types.

**FIGURE 6.**
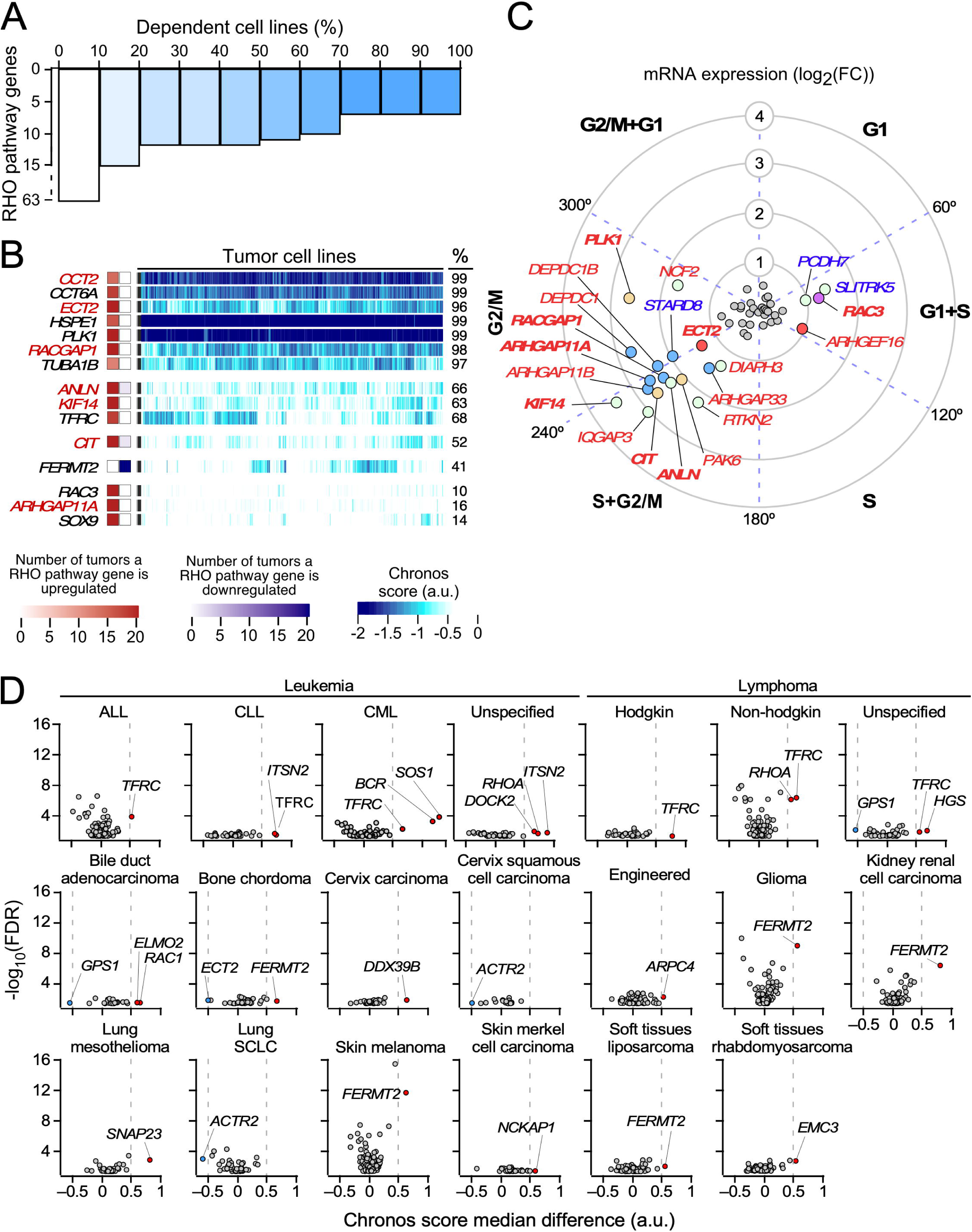
Cancer cell line dependence on the expression of pan-cancer deregulated RHO pathway genes. **(A)** Bar plot representing the percentage of cancer cell lines associated with a dependency score < -0.5 (Chronos score) when a pan-cancer deregulated RHO pathway gene is knocked-out. **(B)** Heatmap showing the Chronos scores associated with pan-cancer deregulated RHO pathway genes in the 1,233 interrogated cancer cell lines. Only genes associated with a score < –0.5 in more than 10% of the analyzed cell lines are represented. DEG, differentially expressed gene. Squares on the left show the differential expression status of these genes across TCGA cohorts (red, upregulated; blue, downregulated). **(C)** Association between the cell cycles stages of HEK 293T cells and the expression of pan-cancer deregulated *RHO* pathway genes. In bold, genes essential for the proliferation of more than 10% of the analyzed cancer cell lines. **(D)** Vulnerabilities of cancer cell lines representative of indicated cancer types (top) to the depletion of the expression of RHO pathway genes. Y axis values represent the - log10(FDR adjusted *P*) associated with each gene when comparing the distribution of Chronos scores between a subgroup of tumor cells from the same subtype with another subgroup gathering all the remaining cell lines from different subtypes in the study. X axis values represent the difference between both subgroups median Chronos scores for each gene. Genes associated with a |median difference| > 0.5 and a *P* < 0.01 are considered to exert a significant impact on the proliferation of cancer cell lines from a particular tumor subtype. ALL, acute lymphocytic leukemia; CLL, chronic lymphocytic leukemia; CML, chronic myelogenous leukemia; Engineered, genetically engineered cancer cell lines; SCLC, small cell lung cancer.

These patterns of coregulated expression suggested that the transcriptional alterations found in pan-cancer differentially expressed RHO GTPase pathway genes could be caused by their participation in broader, pan-cancer regulatory gene expression networks. To further explore this possibility, we expanded the list of coregulated genes by identifying the genes of the human genome that show positive or negative coregulation with the differentially expressed pan-cancer RHO GEFs and RHO GAPs. We then used that information to build a RHO GTPase-centered, weighted gene lists to carry out gene set enrichment analysis (GSEA) using Gene Ontology (GO) biological processes. The top 100 GO terms per group were ranked by median normalized enrichment score (NES) across tumor types. For RHOA-specific networks, enriched processes included tumor angiogenesis, MAPK signaling, inhibition of apoptosis, tumor growth, and cell migration (**Fig. 5D**, red boxes). For RAC1-specific networks, enriched processes were associated with transcriptional programs involved in membrane projection dynamics, DNA damage responses, and metabolic regulation (**Fig. 5D**, blue boxes). Importantly, 50% of the top GO terms were shared between the RHOA- and RAC1-specific networks (**Fig. 5D**, brown boxes). Together, these data indicate that many cancers engage a coordinated, pan-cancer transcriptional programs that include the modulation of RHO GEF- and RHO GAP-encoding genes.

### Functional impact of the elimination of RHO pathway genes in cancer cell lines

To investigate whether the depletion of RHO pathway genes confers a proliferative disadvantage to tumor cells, we next assessed the dependency of 1,032 cancer cell lines on the expression of RHO pathway genes using data from genome-wide CRISPR/Cas9 essentiality screens [34, 35]. Among the 129 pan-cancer deregulated RHO pathway genes identified in TCGA tumors (**Fig. 3**), the depletion of 63 genes significantly reduced proliferation in at least one cancer cell line (Chronos score < –0.5) (**Fig. 6A**). Notably, 15 of these genes were required for the growth of more than 10% of all tested cell lines, indicating a broad spectrum of dependency (*CCT2*, *CCT6A*, *ECT2*, *HSP1*, *PLK1*, *RACGAP1*, *TUBA1B*, *ANLN*, *KIF14*, *TFRC*, *CIT*, *FERMT2*, *RAC3*, *ARHGAP11A*, *SOX9*) (**Fig. 6B**). 4 of those 15 genes were detected in our SCNV studies (*ANLN*, *ECT2*, *KIF14*, *TFRC*) (**Fig. 4D**). The expression of 7 of those 15 genes was also strongly associated with cell cycle progression, particularly during the G1+S (RAC3), S+G_2_/M (*ANLN*, *ARHGAP11A*, *CIT*, *ECT2*, *KIF14*), and G_2_/M (*PLK1*, *RACGAP1*) phases based on previous studies on cell cycle-regulated genes [65] (**Fig. 6C**, genes in bold and red font). These results suggest that their depletion disrupts cell cycle progression, thus explaining the widespread growth inhibition observed upon their knockout. Our data, however, support the idea of a more general involvement in many cell types.

We next asked whether RHO pathway gene dependencies vary in a tumor type– specific manner. To this end, we stratified cell lines based on tissue of origin and compared the distribution of Chronos scores for each RHO pathway gene within each tumor group against all other cancer types combined. This analysis produced two values per gene-cancer type pair: **(i)** the difference in median Chronos scores, and **(ii)** a corresponding FDR-adjusted *P* value from a two-group comparison. Genes showing a median difference > 0.5 and FDR-adjusted *P* < 0.01 were considered to exhibit significant tumor-specific vulnerability. This approach revealed multiple lineage-specific dependencies. In hematopoietic malignancies, *TFRC*, a pan-cancer upregulated RHO pathway gene (**Fig. 3B**), was essential in all leukemia and lymphoma-derived lines (**Fig. 6D**). Chronic lymphocytic leukemia (CLL) cells showed further dependency on *ITSN2* (**Fig. 6D**); chronic myeloid leukemia (CML) cell lines showed also dependency on *SOS1* and *BCR* ; and non-Hodkin lymphomas on *RHOA* (**Fig. 6D**). The role of BCR in these cells must be due to the downmodulation of the oncogenic BCR-ABL1 fusion protein typical of this tumor type rather than to the intrinsic function of BCR. Unspecified leukemia cell lines were dependent on *DOCK2*, *ITSN2* and *RHOA* whereas unspecified lymphoma cells were *HGS*- and *GPS1*-dependent (**Fig. 6D**). In solid tumors, *FERMT2* was important for the proliferation of cell lines from bone chordoma (together with *ECT2*), glioma, kidney renal cell carcinoma, skin melanoma, and soft tissues sarcoma (**Fig. 6D**). *ELMO2*, *RAC1* and *GPS1* were important for bile duct adenocarcinoma; *DDX39B* for cervix carcinoma, *ACTR2* for both cervix squamous cell carcinoma and small cell lung cancer; *SNAP23* for lung mesothelioma; *NCKAP1* for skin Merkel cell carcinoma; and *EMC3* for soft tissues rhabdomyosarcoma (**Fig. 6D**). Interestingly, we observed that the depletion of some of genes led to higher proliferation rates in some cell lines (*ACTR2*, *ECT2*, *GPS1*) (**Fig. 6D**, blue circles). Together, these analyses underscore the functional relevance of the deregulation of some RHO GTPase pathway genes for the potential fitness of specific cancer types.

## CONCLUDING REMARKS

In this study we have integrated genomic, transcriptomic, and functional dependency data from large-scale public resources to generate the first comprehensive dissection of the alterations found in the RHO GTPase signaling landscape across human cancers. As inferred from previous analyses [1, 3, 4], our findings indicate that the contribution of genetic alterations (mutations, SNCVs) is rather limited. Despite this, we have found a small, although significant number of genes that contain positively selected mutations, hotspot mutations and/or SNCVs in specific TCGA cancer subtypes (**Supplementary Table S3**). Those genes encode 4 GTPases (*RAC1*, *RHOA*, *RHOB*, *RND1*), 8 DH domain-containing proteins (*ARHGEF15*, *ECT2*, *FGD5*, *MCF2*, *PLEKHG2*, *PLEKHG4*, *PLEKHG4B*, *PLEKHG7*), a DOCK domain-containing protein (*DOCK1*), 13 RHO GAP domain-containing proteins (*ARAP2*, *ARHGAP1*, *ARHGAP9*, *ARHGAP29*, *ARHGAP30*, *ARHGAP35*, *ARHGAP39*, *CHN2*, *DEPDC1*, *DEPDC1B*, *PIK3R1, PIK3R2*, *STARD13*), 10 kinases (*CDC42BPA*, *MAP3K1*, *MYLK*, *PAK6*, *PIK3CA*, *PKN2*, *PLK1*, *RPS6KB1*, *STK10*, *TNK1*), and 32 proximal or distal signaling elements many of which encode cytoskeletal or cell polarity components (**Supplementary Table S3**). In some cases, two types of genetic alterations are found in the same gene (e.g., positively selected and hotspot mutations as in the case of *RAC1*, *RHOA*, *RHOB*, *ARHGAP35*, *PIK3R1*, *PIK3CA*, *PLK1*, and *ACTB*; mutations and gene copy number losses as the case of *MYH1*1, indicating that they are probably the most likely candidates to act as oncogenic drivers. These latter genes (underlined) and other loci (not underlined) can also undergo genetic alterations and changes in expression (*<u>RHOB</u>*, *ARHGAP39*, *DEPDC1B*, *STARD13*, *ARHGEF15*, *ECT2*, *FGD5*, *PLEKHG*2, *PLEKHG4*, *PLEKHG4B*, *DOCK1*, *MAP3K1*, *MYLK*, *PAK6*, *<u>PIK3CA</u>*, *<u>PLK1</u>*, *ACTC1*, *ACTR3B*, *ANLN*, *ARL13B*, *FAM65C*, *IQGAP3*, *KIF4*, *MTMR1*, *<u>MYH11</u>*, *NHS*, *POTEE*, *RHPN1*, *SH3RF2*, *SLC4A7*, *SLITRK3*, *SOX9*, and *TFRC*) (**Supplementary Table S3**), further indicating that such alterations might have functional relevance. A minority of genes having such genetic alterations are also important for the proliferation of a larger subset of cancer cell lines (*RAC1*, *ECT2*, *PLK1*, *ANLN*, *KIF14*, *SOX9*) or for cancer cell lines representative of specific tumor subtypes (*RAC1*, *RHOA*, *TFRC*) according to CRISPR-CAS9 gene-editing-based screenings.

Despite the foregoing data, our results indicate as anticipated [1, 3, 4] that the most common deregulation found for RHO GTPase pathway genes is by far that associated with changes in transcript expression. In some cases, such changes affect genes that do not undergo statistically significant genetic alterations in the TCGA cancer subtypes interrogated in this study. Some of those genes also have been found to be important for the proliferation of either a large number (*RAC3*, *ARHGAP11A*, *RACGAP1*, *CIT*, *CCT2*, *CCT6A*, *FERMT2*, *HSPE1*, *TUBA1B*) or a specific number (*BCR*, *GPS1*, *HGS*) of cancer cell lines (**Fig. 6**). This indicates that the changes of gene expression of these genes are likely relevant for oncogenesis. Our study also suggests that the deregulation of the expression of most RHO GTPase pathway genes cannot be interpreted as the consequence of just their intrinsic or individual functional impact on cancer cells but, rather, by being components of large transcriptional programs that become activated in cancer cells (**Fig. 5**). Thus, rather than being drivers *per se*, it is likely that these proteins will mostly work as ancillary factors that participate of large-scale biological processes regulated by yet unidentified transcriptional factors.

Our results also underscore the complexity of the regulation of RHO GTPases in cancer. For example, the discordance between canonical expectations and observed expression patterns found among many RHO GEFs and RHO GAPs suggests the existence of context-dependent, non-canonical functions that warrant further mechanistic exploration. Alternatively, as discussed above, it can merely reflect bystander events or the enrichment of a cell type that developed the tumor and that had a different expression pattern of those transcripts when compared to the rest of cell types of the tissue where the disease has originated from.

While our study provides a comprehensive analysis to date of RHO signaling dysregulation in cancer, the readers must be aware that several limitations remain. Firstly, it is worth mentioning that the functional impact of many identified mutations (especially those that are not positively selected or not found in hotspots) and differential expression patterns found across cancers in our work remains to be experimentally validated. Secondly, it must be underscored that the deregulations observed at the transcript level do not always have to match the expression or activity levels of the encoded proteins. Thirdly, changes in mRNA and protein expression might not always correlate with a functional impact at the cellular level due to the activation of buffering signaling events. Our analyses will not pick up genes encoding signaling components of RHO GTPase-regulated pathways whose activity is modulated by upstream oncogenic signals rather than by mutations or gene expression changes. Finally, it is also worth indicating that the functional screenings may yield incorrect information in the case of sgRNAs that do not property inactivate the expected target gene. To overcome these problems, our studies must be complemented with additional phosphoproteomic, proteomic, and activity-based profiling analysis in the future to adequately identify the full spectrum of mechanism by which RHO GTPase-regulated pathways impact on cancer cell biology and pathophysiology. Regardless of these caveats, our study has generated a comprehensive catalogue of genetic and non-genetic mechanisms affecting RHO GTPase pathway genes in cancer. This roadmap will be relevant to guide future efforts aimed at targeting RHO GTPase-regulated pathways in oncological malignancies.

## EXPERIMENTAL PROCEDURES

### Curation of a list of genes involved in the RHO GTPase pathways

To elaborate a list of genes involved in the RHO GTPase-dependent signaling pathways we followed three criteria of inclusion: (i) genes encoding the 23 RHO GTPases; (ii) genes encoding proteins involved in the regulation of the RHO GTPase cycle; (iii) genes encoding proteins that have been described as either proximal or distal effectors in previous publications; (iv) genes encoding proteins that have been described in the proximal interactome of RHO GTPases [36].

### Somatic mutation analyses

TCGA mutation annotation format (MAF) files generated by MuTect2 variant caller pipeline [66] were downloaded from the Genomic Data Commons (GDC) database using the TCGAbiolinks R package [67]. The TCGA cohorts under analysis were those indicated in **Supplementary Table** 2. Then, we calculated the mutational burden in samples using the maftools R package [68]. Those samples with a number of mutations 4.5 times larger than the average number of somatic mutations found in the respective analyzed cohort were classified as “hypermutated” and removed from the analysis. From filtered MAF files, we calculated RHO GTPase pathway mutational load and prevalence as well as the background distributions of genes mutational load and prevalence in each cohort. To that end, we applied a random sampling approach to create 1,000 lists with the same number of genes as the list of RHO pathway genes (484). These new lists are similar in terms of gene length distribution to the list of RHO pathway genes. This was done to avoid biases in our results from the fact that longer genes often present larger mutational loads in cancer. The normality of each background distribution was assessed using fitdistrplus R package (https://cran.r-project.org/web/packages/fitdistrplus/index.html). RHO GTPase pathway and background distribution values were then transformed into Z-scores. Comparisons were finally performed by using the Poisson test. *P* values were further adjusted using the false discovery rate (FDR) method. Significant threshold was set on an FDR adjusted *P* < 0.05.

### Analysis of putative oncogenic mutations in RHO pathway genes

We applied two bioinformatic methods on the filtered MAF files. Firstly, we measured positive selection signals on RHO pathway genes mutations using the dNdScv R package [69]. Genes associated with a qglobal value < 0.01 were considered to harbor mutations under significant positive selection. Genes qmiss_cv and qtrunc_cv values were then used, respectively, to differentiate whether missense or truncating mutations were significantly selected on them. Secondly, we applied the OncodriveCLUSTL algorithm [70] to identify clustered mutations (“mutational spots”) on *RHO* pathway genes. This method was run in each cohort with the following parameters: “smooth windows” of 11, a “simulation window” of 31, a “mutation cluster cutoff” of 2, 1,000 simulations and the simulation mode as “mutation_centered”. Finally, we calculated the proportion of a gene’s missense and truncating mutations harbored at these hotspots.

### Differential expression analyses on TCGA datasets

TCGA RNAseq raw counts data derived from Illumina HiSeq platform were downloaded from the Genomic Data Commons (GDC) server, processed, and normalized using the TCGAbiolinks R package [71]. Firstly, we performed an array-array intensity correlation (AAIC) analysis to remove outlier samples with an associated whole-transcriptome Pearson correlation value lower than 0.6. Then, genes average expression was calculated and those within the lowest expressed quartile were filtered out. Finally, the resulting matrix of counts was further normalized following the EDAseq R package pipeline [72] in two steps: (i) a “within-lane” normalization step to reduce counts dependence on genes GC content; and (ii) a “between-lane” normalization step to reduce their dependence on inter-samples sources of variation. Once normalized, we applied the edgeR differential expression analysis (DEA) pipeline [73], integrated in the TCGAbiolinks R package. Genes with an associated |log_2_(FC)| value > 1 and an FDR adjusted *P < 0.01* were considered differentially expressed.

### Integration of somatic copy number and mRNA expression data

To carry out these analyses, we developed a new R-based method compiled in the CiberAMP R package [64]. Briefly, the data of somatic copy number variations of RHO pathway genes were downloaded from the Broad’s Institute Firebrowser using TCGAbiolinks R package. We acquired the “thresholded-by-gene” file from the latest GISTIC2.0 run from each cohort [74]. Then, tumor samples were classified according to RHO pathway gene genotype as amplified, deleted or diploid. We analyzed separately deep and shallow copy number altered samples to associate them with the transcriptional deregulation of *RHO* pathway genes in each TCGA cohort. Genes associated with a |log_2_(FC)| value > 1 and FDR adjusted *P* < 0.01 were considered differentially expressed.

Next, we calculated the co-occurrence between the somatic copy number variations of *RHO* pathway genes and known oncogenes in each cohort. To that end, GISTIC2.0 “thresholded-by-gene” file information [75] was encoded into two binary matrices. On the one hand, in the first matrix values of 1 encode a deep copy number variation (amplification or deletion) of a given gene in a given tumor sample, and values of 0 encode a diploid state. On the other hand, in the second matrix values of 1 encode a shallow copy number variation (amplification or deletion) of a given gene in a given sample, and values of 0 a diploid state. Then, we calculated which genes are significantly co-amplified or co-deleted in each cohort. To that end, for every pair of genes within each matrix, we calculated the co-occurrence *P* between their copy number variations using Fisher’s exact test. Two genes were considered significantly co-amplified or co-deleted when associated with an adjusted *P* value < 0.05.

### Gene set enrichment analyses

For these analyses, we followed the steps described in [76]. Briefly, we computed pairwise Spearman correlation coefficients between the pan-cancer up- and down-regulated RHO GEFs and RHO GAPs with specific activity for RAC1 and RHOA GTPases, and every other transcript in each cohort. The ranked gene list based on decreasing order of Spearman coefficients was used as input to the gseGO function of clusterProfiler R package [77] to evaluate genes enrichment on biological processes using 10,000 permutations, a maximum gene set size of 800 and a *P* cutoff of 0.01. Finally, we selected top 100 correlated biological processes from each cohort according to their associated normalized enrichment score (NES).

### Genetic dependence of cancer cell lines on RHO GTPase pathway genes

1,032 tumor cell lines genetic dependency scores from CRISPR/Cas9 genome-wide screens (DepMap 21Q3 Public+Score, Chronos score) and their annotated somatic mutations were downloaded from the Dependency Map (DepMap) database [34, 35]. We considered as pan-cancer essential all those genes associated with a Chronos score < –0.5 in more than 10% of the analyzed tumor cell lines. Then, we compared genes associated Chronos scores between tumor cell lines lineages. To that end, we classified cell lines according to their tumor of origin. For each resulting subset of cell lines, we calculated their median Chronos score. This score was then compared to the median Chronos score of all the remaining cell lines using parametric (Student’s *t* test) or non-parametric (Welch’s test) statistical tests. The resulting *P* were further adjusted using the false discovery rate (FDR) method. A comparison was considered statistically significant when associated with an FDR adjusted *P* < 0.01.

### Correlation between the expression of RHO pathway gene expression and the cell cycle stages

To find out associations between *RHO* pathway genes expression and the cell cycle of tumor cells, we selected a dataset in which HEK 293T cells were transcriptionally profiled at different stages. The associations between the expression of *RHO* pathway genes and cell cycle stages was obtained from [65].

## MATERIALS AVAILABILITY

All relevant data are available from the corresponding author upon reasonable request. A Materials Transfer Agreement could be required in the case of potential commercial applications.

## AUTHOR CONTRIBUTIONS

R.F. conducted the analyses and designed the figures. L.F.L.-M. and V.Q. helped in the design of *in silico* strategies and the analysis of the data obtained. X.R.B. conceived the work, analyzed data, wrote the manuscript, and carried out the final editing of figures.

## ACKNOWLEDGEMENTS

The X.R.B.’s project leading to these results has received funding from the Spanish Association against Cancer (GC16173472GARC), the Castilla-León government (CSI018P23), grants funded by MCIN/AEI/10.13039/501100011033/ plus the European Research Development Fund «A way of making Europe» of the European Union (PID2021-122666OB-I00, PDC2022-133027-I00, PLEC2022-009217), «la Caixa» Banking Foundation (HR20-00164), the «Programa Excelencia» of the Fundación Científica AECC 2022 (EPAEC222641CICS), and the «Escalera de Excelencia» of the Education Ministry of the Castilla y León autonomous government plus the European Research Development Fund (CLU-2023-2-01).

