## Supplemental Information for "PAN-CANCER ANALYSES IDENTIFY ONCOGENIC DRIVERS, EXPRESSION SIGNATURES, AND THERAPEUTIC VULNERABILITIES IN RHO GTPase PATHWAY GENES"

by

R. Fernández *et al*.

This PDF file includes:

Supplementary Figures 1 to 4 and legends

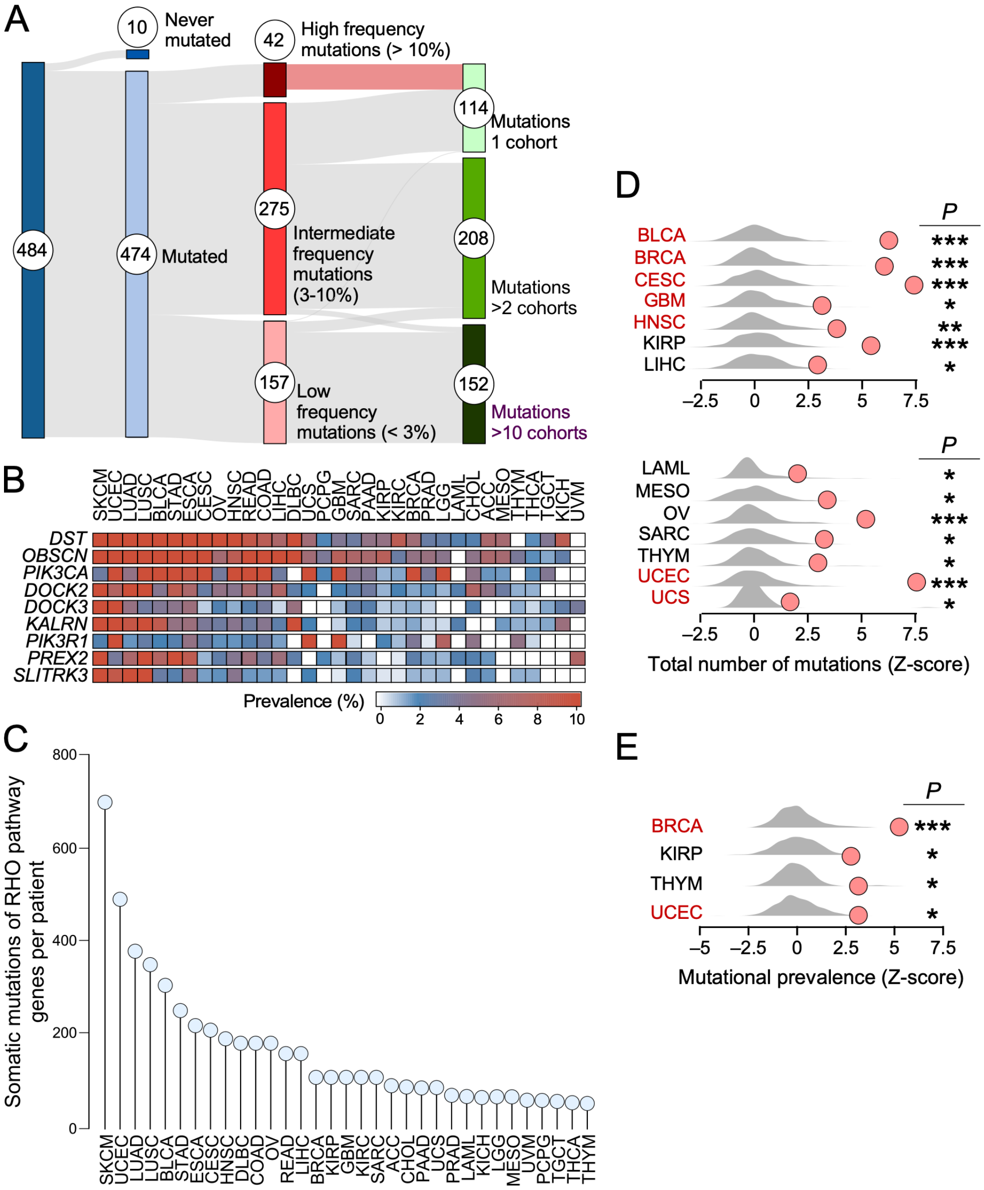


**SUPPLEMENTARY FIGURE 1. Somatic mutational landscape of *RHO* pathway genes in TCGA tumors**

**(A)** Sankey diagram with a summary of the information obtained from the analysis of the somatic mutations found on *RHO* pathway genes across TCGA tumors.

**(B)** Heatmap representing the mutational prevalence of 9 RHO pathway genes mutated in more than one cohort and in at least 10% of the patients. TCGA tumor abbreviations are those shown in **Supplementary Table 2**.

**(C)** Somatic mutation burden of RHO pathway genes in indicated tumor types (bottom).

**(D and E)** Results from the random sampling analysis of the mutational load (D) and prevalence (E) of RHO pathway genes across TCGA tumors comparing RHO pathway genes associated values (red dots) with an expected background distribution (grey shapes). *, $P < 0.05$; **, $P < 0.005$; ***, $P < 0.0005$ (Poisson test). Only significant results are represented.

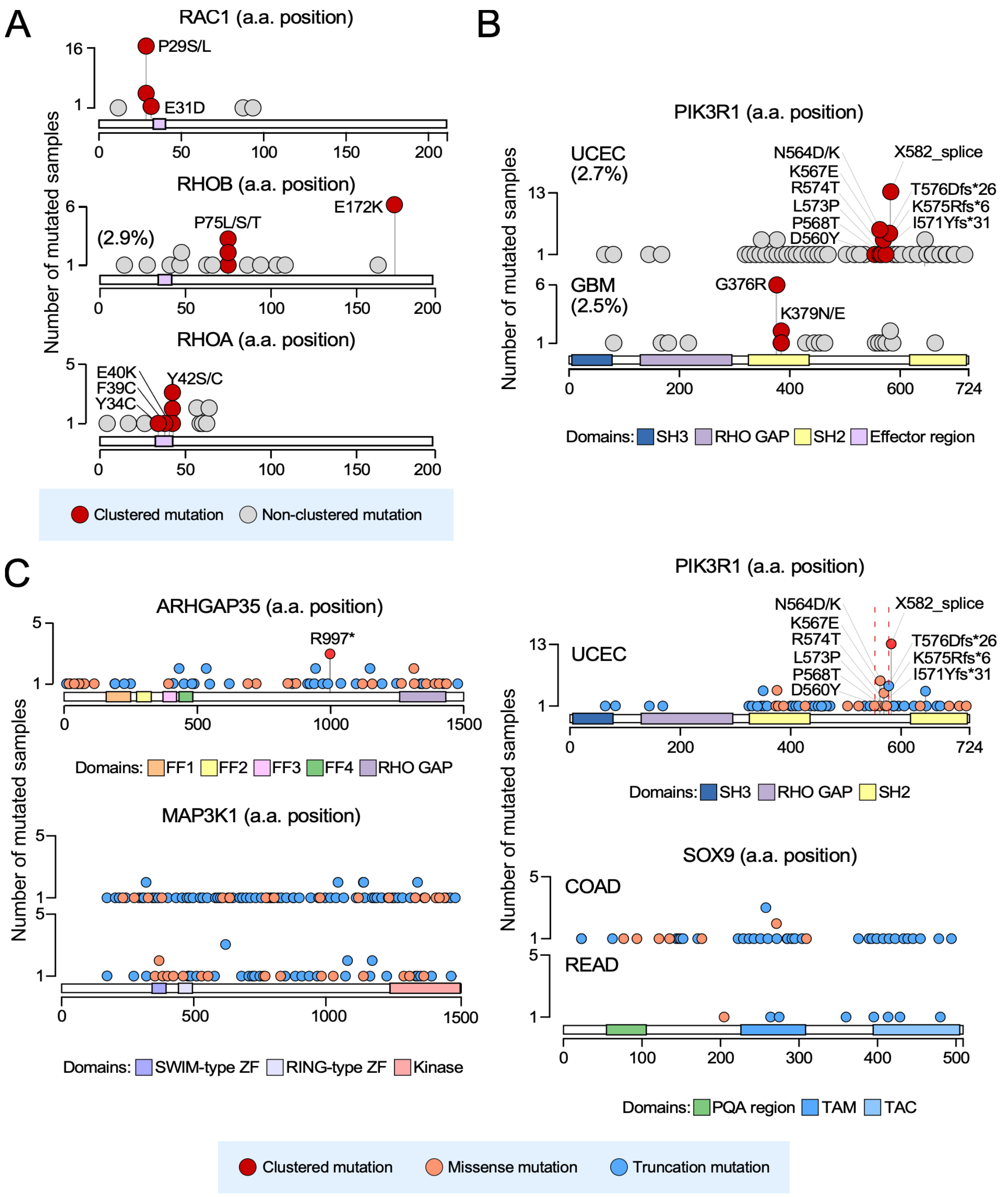


**SUPPLEMENTARY FIGURE 2. Examples of hotspot mutations found in RHO pathway genes**

**(A** and **B)** Needle plots of top prevalent putative oncogenic mutations (red dots) identified among RHO pathway genes in TCGA tumors. a.a, amino acid; *, STOP codon; fs*, frameshift mutation; _splice, splicing mutation

**(C)** Needle plots of somatic mutations on RHO pathway genes with truncating alterations under significant positive selection. Dashed lines depict the limits of the missense-enriched mutational cluster found on *PIK3R1* in UCEC tumors (top panel on the right).

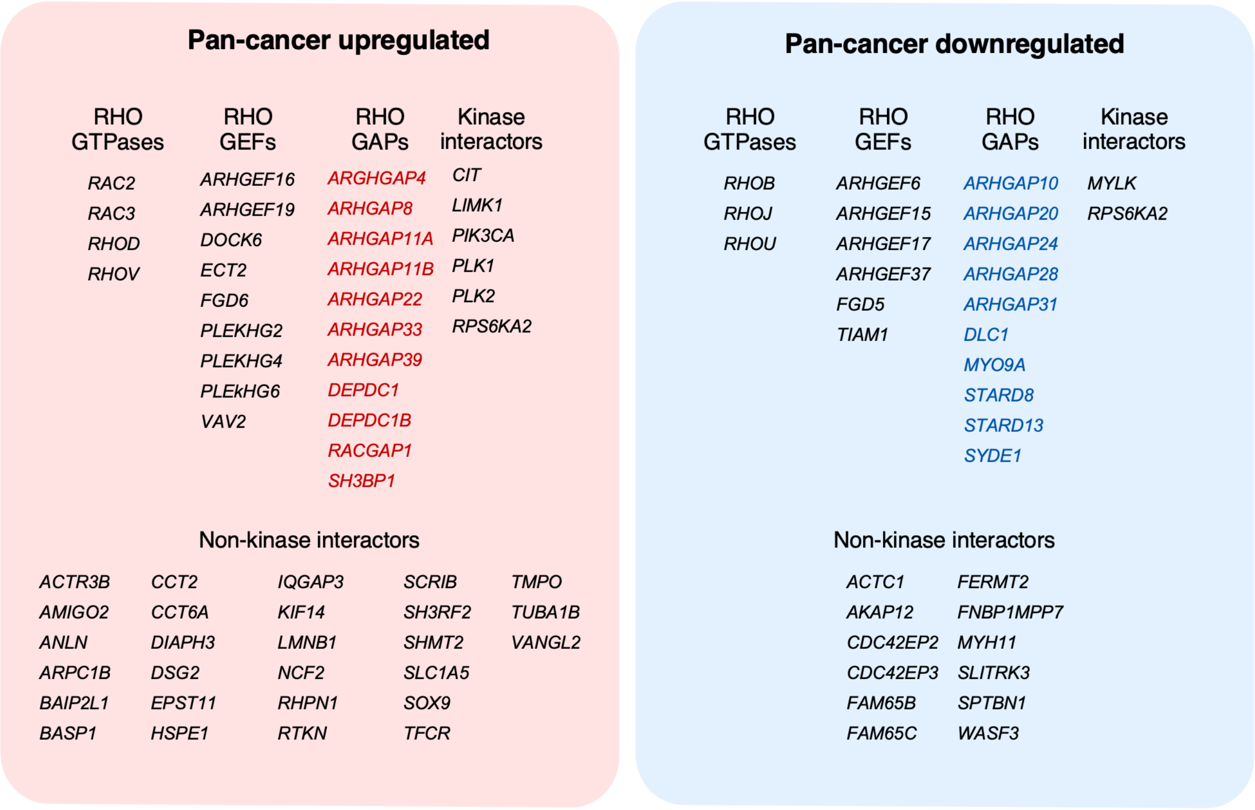


**SUPPLEMENTARY FIGURE 3. RHO pathway genes deregulated in pan-cancer data**

We show RHO pathway genes that are upregulated (red box) and downregulated (blue box) in more than 25% of interrogated TCGA tumor cohorts. We also indicate RHO GAPs (red font) and RHO GEFs (blue font) with expression patterns opposite to those expected according to their canonical functions.

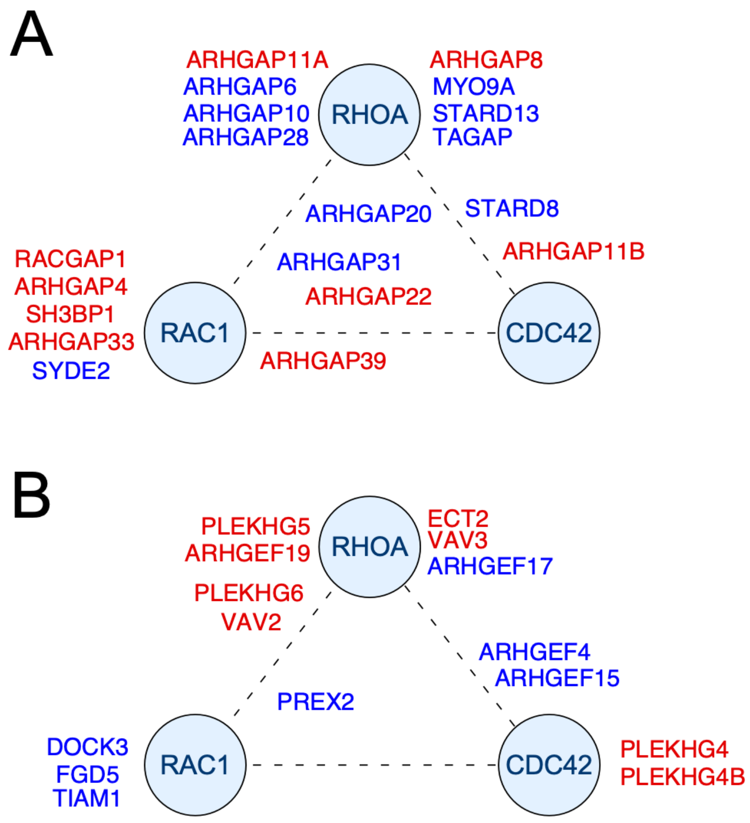


**SUPPLEMENTARY FIGURE 4. Functional analysis of RHO GAPs and GEFs with deregulated expression in pan-cancer data**

**(A and B)** Ternary plots showing the catalytic affinity of pan-cancer upregulated (red gene symbols) and downregulated (blue gene symbols) RHO GAPs (A) and RHO GEFs (B) towards the three main RHO subfamily GTPases (CDC42, RAC1, RHOA). The affinity for a given GTPase is represented by the proximity to its corresponding coordinates. See further details in main text.
