## Supplementary Table 1 for "PAN-CANCER ANALYSES IDENTIFY ONCOGENIC DRIVERS, EXPRESSION SIGNATURES, AND THERAPEUTIC VULNERABILITIES IN RHO GTPase PATHWAY GENES"

### Supplementary Table S1. List of RHO GTPase pathway genes used in this study.

| Symbol | Subgroup | Chr | Band | Start | End |
|---|---|---|---|---|---|
| *CDC42* | RHO GTPase | 1 | p36.12 | 22025511 | 22101360 |
| *RAC1* | RHO GTPase | 7 | p22.1 | 6374527 | 6403967 |
| *RAC2* | RHO GTPase | 22 | q13.1 | 37225270 | 37244448 |
| *RAC3* | RHO GTPase | 17 | q25.3 | 82031678 | 82034204 |
| *RHOA* | RHO GTPase | 3 | p21.31 | 49359139 | 49412998 |
| *RHOB* | RHO GTPase | 2 | p24.1 | 20447074 | 20449440 |
| *RHOBTB1* | RHO GTPase | 10 | q21.2 | 60869438 | 61001440 |
| *RHOBTB2* | RHO GTPase | 8 | p21.3 | 22987417 | 23020199 |
| *RHOBTB3* | RHO GTPase | 5 | q15 | 95713522 | 95824383 |
| *RHOC* | RHO GTPase | 1 | p13.2 | 112701127 | 112707434 |
| *RHOD* | RHO GTPase | 11 | q13.2 | 67056847 | 67072017 |
| *RHOF* | RHO GTPase | 12 | q24.31 | 121777754 | 121803403 |
| *RHOG* | RHO GTPase | 11 | p15.4 | 3826978 | 3840959 |
| *RHOH* | RHO GTPase | 4 | p14 | 40191053 | 40246967 |
| *RHOJ* | RHO GTPase | 14 | q23.2 | 63204114 | 63293508 |
| *RHOQ* | RHO GTPase | 2 | p21 | 46541806 | 46584688 |
| *RHOT1* | RHO GTPase | 17 | q11.2 | 32142454 | 32253374 |
| *RHOT2* | RHO GTPase | 16 | p13.3 | 668105 | 674174 |
| *RHOU* | RHO GTPase | 1 | q42.13 | 228735479 | 228746664 |
| *RHOV* | RHO GTPase | 15 | q15.1 | 40872214 | 40874234 |
| *RND1* | RHO GTPase | 12 | q13.12 | 48857145 | 48865870 |
| *RND2* | RHO GTPase | 17 | q21.31 | 43025231 | 43032041 |
| *RND3* | RHO GTPase | 2 | q23.3 | 150468195 | 150539011 |

| Symbol | Subgroup | Chr | Band | Start | End |
|---|---|---|---|---|---|
| *ABR* | RHO GAP and GEF (DH) | 17 | p13.3 | 1003519 | 1229738 |
| *AKAP13* | RHO GEF (DH) | 15 | q25.3 | 85380571 | 85749358 |
| *ALS2* | RHO GEF (DH) | 2 | q33.1 | 201700267 | 201782112 |
| *ARHGEF1* | RHO GEF (DH) | 19 | q13.2 | 41883173 | 41930150 |
| *ARHGEF2* | RHO GEF (DH) | 1 | q22 | 155946851 | 156007070 |
| *ARHGEF3* | RHO GEF (DH) | 3 | p14.3 | 56727418 | 57079329 |
| *ARHGEF4* | RHO GEF (DH) | 2 | q21.1 | 130836914 | 131047263 |
| *ARHGEF5* | RHO GEF (DH) | 7 | q35 | 144355288 | 144380632 |
| *ARHGEF6* | RHO GEF (DH) | X | q26.3 | 136665547 | 136780932 |
| *ARHGEF7* | RHO GEF (DH) | 13 | q34 | 111114559 | 111305737 |
| *ARHGEF9* | RHO GEF (DH) | X | q11.2 | 63634967 | 63809274 |
| *ARHGEF10* | RHO GEF (DH) | 8 | p23.3 | 1823926 | 1958641 |
| *ARHGEF10L* | RHO GEF (DH) | 1 | p36.13 | 17539698 | 17697874 |
| *ARHGEF11* | RHO GEF (DH) | 1 | q23.1 | 156934840 | 157045742 |
| *ARHGEF12* | RHO GEF (DH) | 11 | q23.3 | 120336413 | 120489937 |
| *ARHGEF15* | RHO GEF (DH) | 17 | p13.1 | 8310241 | 8322514 |
| *ARHGEF16* | RHO GEF (DH) | 1 | p36.32 | 3454665 | 3481113 |
| *ARHGEF17* | RHO GEF (DH) | 11 | q13.4 | 73308276 | 73369388 |
| *ARHGEF18* | RHO GEF (DH) | 19 | p13.2 | 7348937 | 7472485 |
| *ARHGEF19* | RHO GEF (DH) | 1 | p36.13 | 16197854 | 16212652 |
| *ARHGEF25* | RHO GEF (DH) | 12 | q13.3 | 57610180 | 57617245 |
| *ARHGEF26* | RHO GEF (DH) | 3 | q25.2 | 154121003 | 154257827 |
| *ARHGEF28* | RHO GEF (DH) | 5 | q13.2 | 73626158 | 73941993 |
| *ARHGEF33* | RHO GEF (DH) | 2 | p22.1 | 38889875 | 38975454 |
| *ARHGEF35* | RHO GEF (DH) | 7 | q35 | 144186083 | 144195833 |
| *ARHGEF37* | RHO GEF (DH) | 5 | q32 | 149551947 | 149634968 |
| *ARHGEF38* | RHO GEF (DH) | 4 | q24 | 105552620 | 105708093 |
| *ARHGEF39* | RHO GEF (DH) | 9 | p13.3 | 35658875 | 35675866 |
| *ARHGEF40* | RHO GEF (DH) | 14 | q11.2 | 21070273 | 21090248 |
| *BCR* | RHO GEF (DH) and GAP | 22 | q11.23 | 23179704 | 23318037 |
| *DNMBP* | RHO GEF (DH) | 10 | q24.2 | 99875577 | 100009947 |
| *ECT2* | RHO GEF (DH) | 3 | q26.31 | 172750682 | 172821474 |
| *ECT2L* | RHO GEF (DH) | 6 | q24.1 | 138795911 | 138904070 |
| *FARP1* | RHO GEF (DH) | 13 | q32.2 | 98142562 | 98455176 |
| *FARP2* | RHO GEF (DH) | 2 | q37.3 | 241356285 | 241494841 |
| *FGD1* | RHO GEF (DH) | X | p11.22 | 54445454 | 54496234 |
| *FGD2* | RHO GEF (DH) | 6 | p21.2 | 37005646 | 37029069 |
| *FGD3* | RHO GEF (DH) | 9 | q22.31 | 92947523 | 93036236 |

| Symbol | Subgroup | Chr | Band | Start | End |
|---|---|---|---|---|---|
| *FGD4* | RHO GEF (DH) | 12 | p11.21 | 32399558 | 32646050 |
| *FGD5* | RHO GEF (DH) | 3 | p25.1 | 14810853 | 14934571 |
| *FGD6* | RHO GEF (DH) | 12 | q22 | 95076749 | 95217482 |
| *ITSN1* | RHO GEF (DH) | 21 | q22.11 | 33642400 | 33899861 |
| *ITSN2* | RHO GEF (DH) | 2 | p23.3 | 24202864 | 24360536 |
| *KALRN* | RHO GEF (DH) | 3 | q21.1 | 124033369 | 124726325 |
| *MCF2* | RHO GEF (DH) | X | q27.1 | 139581770 | 139708227 |
| *MCF2L* | RHO GEF (DH) | 13 | q34 | 112894378 | 113099742 |
| *MCF2L2* | RHO GEF (DH) | 3 | q27.1 | 183178041 | 183428778 |
| *NET1* | RHO GEF (DH) | 10 | p15.1 | 5412557 | 5459056 |
| *NGEF* | RHO GEF (DH) | 2 | q37.1 | 232878701 | 233013256 |
| *OBSCN* | RHO GEF (DH) | 1 | q42.13 | 228208044 | 228378876 |
| *PLEKHG1* | RHO GEF (DH) | 6 | q25.1 | 150599883 | 150843665 |
| *PLEKHG2* | RHO GEF (DH) | 19 | q13.2 | 39412669 | 39428415 |
| *PLEKHG3* | RHO GEF (DH) | 14 | q23.3 | 64704102 | 64750249 |
| *PLEKHG4* | RHO GEF (DH) | 16 | q22.1 | 67277510 | 67289499 |
| *PLEKHG4B* | RHO GEF (DH) | 5 | p15.33 | 92151 | 189972 |
| *PLEKHG5* | RHO GEF (DH) | 1 | p36.31 | 6467122 | 6520074 |
| *PLEKHG6* | RHO GEF (DH) | 12 | p13.31 | 6310436 | 6328506 |
| *PLEKHG7* | RHO GEF (DH) | 12 | q22 | 92702843 | 92772455 |
| *PREX1* | RHO GEF (DH) | 20 | q13.13 | 48624252 | 48827999 |
| *PREX2* | RHO GEF (DH) | 8 | q13.2 | 67952046 | 68237032 |
| *SPATA13* | RHO GEF (DH) | 13 | q12.12 | 23979805 | 24307074 |
| *TIAM1* | RHO GEF (DH) | 21 | q22.11 | 31118416 | 31559977 |
| *TIAM2* | RHO GEF (DH) | 6 | q25.2 | 154832697 | 155257723 |
| *TRIO* | RHO GEF (DH) | 5 | p15.2 | 14143342 | 14532128 |
| *VAV1* | RHO GEF (DH) | 19 | p13.3 | 6772708 | 6857366 |
| *VAV2* | RHO GEF (DH) | 9 | q34.2 | 133761894 | 133992604 |
| *VAV3* | RHO GEF (DH) | 1 | p13.3 | 107571161 | 107965180 |
| *RASGRF1* | DH & CDC25 domains | 15 | q25.1 | 78959906 | 79090780 |
| *RASGRF2* | DH & CDC25 domains | 5 | q14.1 | 80960363 | 81230162 |
| *SOS1* | DH & CDC25 domains | 2 | p22.1 | 38981549 | 39124345 |
| *SOS2* | DH & CDC25 domains | 14 | q21.3 | 50117130 | 50231578 |
| *DOCK1* | RHO GEF (DOCK) | 10 | q26.2 | 126905409 | 127452517 |
| *DOCK2* | RHO GEF (DOCK) | 5 | q35.1 | 169637268 | 170083382 |
| *DOCK3* | RHO GEF (DOCK) | 3 | p21.2 | 50674927 | 51384198 |
| *DOCK4* | RHO GEF (DOCK) | 7 | q31.1 | 111726110 | 112206407 |
| *DOCK5* | RHO GEF (DOCK) | 8 | p21.2 | 25184689 | 25418082 |

| Symbol | Subgroup | Chr | Band | Start | End |
|---|---|---|---|---|---|
| *DOCK6* | RHO GEF (DOCK) | 19 | p13.2 | 11199295 | 11262524 |
| *DOCK7* | RHO GEF (DOCK) | 1 | p31.3 | 62454298 | 62688386 |
| *DOCK8* | RHO GEF (DOCK) | 9 | p24.3 | 214854 | 465259 |
| *DOCK9* | RHO GEF (DOCK) | 13 | q32.3 | 98793429 | 99086625 |
| *DOCK10* | RHO GEF (DOCK) | 2 | q36.2 | 224765090 | 225042468 |
| *DOCK11* | RHO GEF (DOCK) | X | q24 | 118495815 | 118686163 |
| *RAP1GDS1* | RHO GEF (armadillo) | 9 | q34.13 | 131576770 | 131740076 |

.../...

| Symbol | Subgroup | Chr | Band | Start | End |
|---|---|---|---|---|---|
| *ABR* | RHO GAP and GEF (DH) | 17 | p13.3 | 1003519 | 1229738 |
| *ARAP1* | RHO GAP domain | 11 | q13.4 | 72685069 | 72793599 |
| *ARAP2* | RHO GAP domain | 4 | p14 | 35948221 | 36244514 |
| *ARAP3* | RHO GAP domain | 5 | q31.3 | 141653401 | 141682230 |
| *ARHGAP1* | RHO GAP domain | 11 | p11.2 | 46677080 | 46700619 |
| *ARHGAP4* | RHO GAP domain | X | q28 | 153907367 | 153934999 |
| *ARHGAP5* | RHO GAP domain | 14 | q12 | 32076114 | 32159728 |
| *ARHGAP6* | RHO GAP domain | X | p22.2 | 11117651 | 11665920 |
| *ARHGAP8* | RHO GAP domain | 22 | q13.31 | 44752558 | 44862788 |
| *ARHGAP9* | RHO GAP domain | 12 | q13.3 | 57472264 | 57488814 |
| *ARHGAP10* | RHO GAP domain | 4 | q31.23 | 147732063 | 148072776 |
| *ARHGAP11A* | RHO GAP domain | 15 | q13.3 | 32615144 | 32639941 |
| *ARHGAP11B* | RHO GAP domain | 15 | q13.2 | 30624494 | 30649529 |
| *ARHGAP12* | RHO GAP domain | 10 | p11.22 | 31805398 | 31928876 |
| *ARHGAP15* | RHO GAP domain | 2 | q22.2 | 143091362 | 143768352 |
| *ARHGAP17* | RHO GAP domain | 16 | p12.1 | 24919389 | 25015666 |
| *ARHGAP18* | RHO GAP domain | 6 | q22.33 | 129576132 | 129710177 |
| *ARHGAP19* | RHO GAP domain | 10 | q24.1 | 97222173 | 97292673 |
| *ARHGAP20* | RHO GAP domain | 11 | q23.1 | 110577042 | 110713189 |
| *ARHGAP21* | RHO GAP domain | 10 | p12.1 | 24583609 | 24723887 |
| *ARHGAP22* | RHO GAP domain | 10 | q11.23 | 48446036 | 48656265 |
| *ARHGAP23* | RHO GAP domain | 17 | q12 | 38419280 | 38512385 |
| *ARHGAP24* | RHO GAP domain | 4 | q21.23 | 85475150 | 86002668 |
| *ARHGAP25* | RHO GAP domain | 2 | p13.3 | 68679601 | 68826833 |
| *ARHGAP26* | RHO GAP domain | 5 | q31.3 | 142770377 | 143229011 |
| *ARHGAP27* | RHO GAP domain | 17 | q21.31 | 45393902 | 45434421 |
| *ARHGAP28* | RHO GAP domain | 18 | p11.31 | 6729716 | 6915716 |
| *ARHGAP29* | RHO GAP domain | 1 | p22.1 | 94148988 | 94275068 |
| *ARHGAP30* | RHO GAP domain | 1 | q23.3 | 161046946 | 161069970 |
| *ARHGAP31* | RHO GAP domain | 3 | q13.32 | 119294383 | 119420714 |
| *ARHGAP32* | RHO GAP domain | 11 | q24.3 | 128965060 | 129279324 |
| *ARHGAP33* | RHO GAP domain | 19 | q13.12 | 35774532 | 35788822 |
| *ARHGAP35* | RHO GAP domain | 19 | q13.32 | 46860997 | 47005077 |
| *ARHGAP36* | RHO GAP domain | X | q26.1 | 131058346 | 131089885 |
| *ARHGAP39* | RHO GAP domain | 8 | q24.3 | 144529179 | 144605816 |
| *ARHGAP40* | RHO GAP domain | 20 | q11.23 | 38601934 | 38651035 |
| *ARHGAP42* | RHO GAP domain | 11 | q22.1 | 100687288 | 100993941 |
| *ARHGAP44* | RHO GAP domain | 17 | p12 | 12789498 | 12991643 |

| Symbol | Subgroup | Chr | Band | Start | End |
|---|---|---|---|---|---|
| *ARHGAP45* | RHO GAP domain | 19 | p13.3 | 1065922 | 1086627 |
| *BCR* | RHO GAP and GEF (DH) | 22 | q11.23 | 23179704 | 23318037 |
| *CHN1* | RHO GAP domain | 2 | q31.1 | 174798809 | 175005381 |
| *CHN2* | RHO GAP domain | 7 | p14.3 | 29146569 | 29514328 |
| *DEPDC1* | RHO GAP domain | 1 | p31.3 | 68474152 | 68497221 |
| *DEPDC1B* | RHO GAP domain | 5 | q12.1 | 60596912 | 60700190 |
| *DLC1* | RHO GAP domain | 8 | p22 | 13083361 | 13604610 |
| *FAM13A* | RHO GAP domain | 4 | q22.1 | 88725955 | 89111398 |
| *FAM13B* | RHO GAP domain | 5 | q31.2 | 137937960 | 138051961 |
| *GMIP* | RHO GAP domain | 19 | p13.11 | 19629476 | 19643657 |
| *MYO9A* | RHO GAP domain | 15 | q23 | 71822291 | 72118577 |
| *MYO9B* | RHO GAP domain | 19 | p13.11 | 17075777 | 17214537 |
| *OCRL* | RHO GAP domain | X | q26.1 | 129539849 | 129592561 |
| *OPHN1* | RHO GAP domain | X | q12 | 67949349 | 68433913 |
| *PIK3R1* | RHO GAP domain | 5 | q13.1 | 68215756 | 68301821 |
| *PIK3R2* | RHO GAP domain | 19 | p13.11 | 18153163 | 18170532 |
| *RACGAP1* | RHO GAP domain | 12 | q13.12 | 49976923 | 50033136 |
| *RALBP1* | RHO GAP domain | 18 | p11.22 | 9475009 | 9538114 |
| *RALGAPA1* | RHO GAP domain | 14 | q13.2 | 35538352 | 35809304 |
| *SH3BP1* | RHO GAP domain | 22 | q13.1 | 37634654 | 37656117 |
| *SRGAP1* | RHO GAP domain | 12 | q14.2 | 63844700 | 64162217 |
| *SRGAP2* | RHO GAP domain | 1 | q32.1 | 206203346 | 206464436 |
| *SRGAP3* | RHO GAP domain | 3 | p25.3 | 8980591 | 9363053 |
| *STARD13* | RHO GAP domain | 13 | q13.1 | 33103137 | 33350630 |
| *STARD8* | RHO GAP domain | X | q13.1 | 68647666 | 68725842 |
| *SYDE1* | RHO GAP domain | 19 | p13.12 | 15107401 | 15114985 |
| *SYDE2* | RHO GAP domain | 1 | p22.3 | 85156889 | 85201016 |
| *TAGAP* | RHO GAP domain | 6 | q25.3 | 159034468 | 159045152 |
| *ARFGAP2* | ARF & RHO GAP domains | 11 | p11.2 | 47164299 | 47177125 |
| *ARFGAP3* | ARF & RHO GAP domains | 22 | q13.2 | 42796502 | 42858106 |

| Symbol | Subgroup | Chr | Band | Start | End |
| --- | --- | --- | --- | --- | --- |
| *ARHGDIA* | RHO GDI | 17 | q25.3 | 81867721 | 81871378 |
| *ARHGDIB* | RHO GDI | 12 | p12.3 | 14942031 | 14961728 |
| *ARHGDIG* | RHO GDI | 16 | p13.3 | 280450 | 283010 |

| Symbol | Subgroup | Type of protein | Chr | Band | Start | End |
|---|---|---|---|---|---|---|
| *CDC42BPA* | Kinase interactor | Direct effector* | 1 | q42.13 | 226989865 | 227318502 |
| *CDC42BPB* | Kinase interactor | Direct effector | 14 | q32.32 | 102932380 | 103057549 |
| *CDC42BPG* | Kinase interactor | Direct effector | 11 | q13.1 | 64823052 | 64844653 |
| *CIT* | Kinase interactor | Direct effector | 12 | q24.23 | 119685791 | 119877320 |
| *CKB* | Kinase interactor | Distal effector | 14 | q32.33 | 103519667 | 103522833 |
| *DGKA* | Kinase interactor | Direct effector | 12 | q13.2 | 55927319 | 55954027 |
| *DGKG* | Kinase interactor | Direct effector | 3 | q27.3 | 186105668 | 186362234 |
| *DGKQ* | Kinase interactor | Direct effector | 4 | p16.3 | 958887 | 986895 |
| *DMPK* | Kinase interactor | Direct effector | 19 | q13.32 | 45769709 | 45782552 |
| *INPP5B* | Kinase interactor | Direct effector | 1 | p34.3 | 37860697 | 37947057 |
| *LIMK1* | Kinase interactor | Distal effector | 7 | q11.23 | 74082933 | 74122525 |
| *LIMK2* | Kinase interactor | Distal effector | 22 | q12.2 | 31212239 | 31280080 |
| *MAP3K1* | Kinase interactor | Direct effector | 5 | q11.2 | 56815549 | 56896152 |
| *MAP3K2* | Kinase interactor | Direct effector | 2 | q14.3 | 127298668 | 127388465 |
| *MAP3K3* | Kinase interactor | Direct effector | 17 | q23.3 | 63622415 | 63696305 |
| *MAP3K4* | Kinase interactor | Direct effector | 6 | q26 | 160991727 | 161117385 |
| *MAP3K5* | Kinase interactor | Direct effector | 6 | q23.3 | 136557046 | 136792477 |
| *MAP3K6* | Kinase interactor | Direct effector | 1 | p36.11 | 27354067 | 27366961 |
| *MAP3K7* | Kinase interactor | Direct effector | 6 | q15 | 90513573 | 90587072 |
| *MAP3K8* | Kinase interactor | Direct effector | 10 | p11.23 | 30434021 | 30461833 |
| *MAP3K9* | Kinase interactor | Direct effector | 14 | q24.2 | 70722526 | 70809534 |
| *MAP3K10* | Kinase interactor | Direct effector | 19 | q13.2 | 40191426 | 40215575 |
| *MAP3K11* | Kinase interactor | Direct effector | 11 | q13.1 | 65597756 | 65615382 |
| *MAP3K12* | Kinase interactor | Direct effector | 12 | q13.13 | 53479669 | 53500063 |
| *MAP3K13* | Kinase interactor | Direct effector | 3 | q27.2 | 185282941 | 185489094 |
| *MAP3K14* | Kinase interactor | Direct effector | 17 | q21.31 | 45263119 | 45317029 |

| Symbol | Subgroup | Type of protein | Chr | Band | Start | End |
|---|---|---|---|---|---|---|
| *MAP3K15* | Kinase interactor | Direct effector | X | p22.12 | 19360056 | 19515508 |
| *MAP3K19* | Kinase interactor | Direct effector | 2 | q21.3 | 134964485 | 135047468 |
| *MAP3K21* | Kinase interactor | Direct effector | 1 | q42.2 | 233327724 | 233385148 |
| *MYLK* | Kinase interactor | Direct effector | 3 | q21.1 | 123610049 | 123884332 |
| *MYLK2* | Kinase interactor | Direct effector | 20 | q11.21 | 31819308 | 31834689 |
| *MYLK3* | Kinase interactor | Direct effector | 16 | q11.2 | 46702282 | 46790407 |
| *MYLK4* | Kinase interactor | Direct effector | 6 | p25.2 | 2663629 | 2750922 |
| *PAK1* | Kinase interactor | Direct effector | 11 | q14.1 | 77322017 | 77474635 |
| *PAK2* | Kinase interactor | Direct effector | 3 | q29 | 196739857 | 196832647 |
| *PAK3* | Kinase interactor | Direct effector | X | q23 | 110944285 | 111227361 |
| *PAK4* | Kinase interactor | Direct effector | 19 | q13.2 | 39125770 | 39182816 |
| *PAK5* | Kinase interactor | Direct effector | 20 | p12.2 | 9518036 | 9819689 |
| *PAK6* | Kinase interactor | Direct effector | 15 | q15.1 | 40217428 | 40277487 |
| *PIK3CA* | Kinase interactor | Direct effector | 3 | q26.32 | 179148114 | 179240093 |
| *PIP5K1C* | Kinase interactor | Direct effector | 19 | p13.3 | 3630183 | 3700468 |
| *PKN1* | Kinase interactor | Direct effector | 19 | p13.12 | 14433053 | 14471867 |
| *PKN2* | Kinase interactor | Direct effector | 1 | p22.2 | 88684222 | 88836255 |
| *PKN3* | Kinase interactor | Direct effector | 9 | q34.11 | 128702503 | 128720916 |
| *PLK1* | Kinase interactor | Distal effector | 16 | p12.2 | 23677656 | 23690367 |
| *PLK2* | Kinase interactor | Distal effector | 5 | q11.2 | 58453982 | 58460139 |
| *PLK3* | Kinase interactor | Distal effector | 1 | p34.1 | 44800377 | 44805990 |
| *PTK2* | Kinase interactor | Distal effector | 8 | q24.3 | 140657900 | 141002216 |
| *PTK2B* | Kinase interactor | Distal effector | 8 | p21.2 | 27311482 | 27459391 |
| *ROCK1* | Kinase interactor | Direct effector | 18 | q11.1 | 20946906 | 21111813 |
| *ROCK2* | Kinase interactor | Direct effector | 2 | p25.1 | 11179759 | 11348330 |

| Symbol | Subgroup | Type of protein | Chr | Band | Start | End |
|---|---|---|---|---|---|---|
| *RPS6KA1* | Kinase interactor | Distal effector | 1 | p36.11 | 26529761 | 26575030 |
| *RPS6KA2* | Kinase interactor | Distal effector | 6 | q27 | 166409364 | 166906451 |
| *RPS6KB1* | Kinase interactor | Distal effector | 17 | q23.1 | 59893046 | 59950574 |
| *RPS6KB2* | Kinase interactor | Distal effector | 11 | q13.2 | 67428460 | 67435401 |
| *SLK* | Kinase interactor | Direct effector | 10 | q24.33 | 103967140 | 104029233 |
| *STK10* | Kinase interactor | Distal effector | 5 | q35.1 | 172042079 | 172188224 |
| *STK38* | Kinase interactor | Distal effector | 6 | p21.31 | 36493892 | 36547479 |
| *TAOK1* | Kinase interactor | Direct effector | 17 | q11.2 | 29390363 | 29551903 |
| *TAOK2* | Kinase interactor | Direct effector | 16 | p11.2 | 29973868 | 29992261 |
| *TAOK3* | Kinase interactor | Direct effector | 12 | q24.23 | 118149801 | 118372907 |
| *TNK1* | Kinase interactor | Direct effector | 17 | p13.1 | 7380534 | 7389774 |
| *TNK2* | Kinase interactor | Direct effector | 3 | q29 | 195863364 | 195911945 |
| *VRK2* | Kinase interactor | Distal effector | 2 | p16.1 | 57907629 | 58159920 |
| *AAAS* | Non-kinase interactor | Proximal interactor | 12 | q13.13 | 53307456 | 53324864 |
| *ABCD3* | Non-kinase interactor | Proximal interactor | 1 | p21.3 | 94418389 | 94518666 |
| *ABI1* | Non-kinase interactor | Proximal interactor | 10 | p12.1 | 26746593 | 26861087 |
| *ABI2* | Non-kinase interactor | Proximal interactor | 2 | q33.2 | 203328280 | 203447728 |
| *ABL2* | Non-kinase interactor | Proximal interactor | 1 | q25.2 | 179099330 | 179229684 |
| *ACBD5* | Non-kinase interactor | Proximal interactor | 10 | p12.1 | 27168135 | 27243046 |
| *ACTB* | Non-kinase interactor | Direct effector | 7 | p22.1 | 5526409 | 5563902 |
| *ACTC1* | Non-kinase interactor | Direct effector | 15 | q14 | 34790230 | 34795549 |
| *ACTG1* | Non-kinase interactor | Direct effector | 17 | q25.3 | 81509413 | 81523847 |
| *ACTN1* | Non-kinase interactor | Direct effector | 14 | q24.1 | 68874128 | 68979440 |
| *ACTR2* | Non-kinase interactor | Direct effector | 2 | p14 | 65227788 | 65271253 |
| *ACTR3* | Non-kinase interactor | Direct effector | 2 | q14.1 | 113890063 | 113962596 |

| Symbol | Subgroup | Type of protein | Chr | Band | Start | End |
|---|---|---|---|---|---|---|
| *ACTR3B* | Non-kinase interactor | Direct effector | 7 | q36.1 | 152759749 | 152855378 |
| *ACTR3C* | Non-kinase interactor | Direct effector | 7 | q36.1 | 150243916 | 150323725 |
| *ADD3* | Non-kinase interactor | Proximal interactor | 10 | q25.1 | 109996368 | 110135565 |
| *AKAP12* | Non-kinase interactor | Proximal interactor | 6 | q25.1 | 151239967 | 151358559 |
| *ALDH3A2* | Non-kinase interactor | Proximal interactor | 17 | p11.2 | 19648136 | 19685760 |
| *AMIGO2* | Non-kinase interactor | Proximal interactor | 12 | q13.11 | 47075707 | 47079959 |
| *ANKLE2* | Non-kinase interactor | Proximal interactor | 12 | q24.33 | 132725503 | 132761832 |
| *ANKRD26* | Non-kinase interactor | Proximal interactor | 10 | p12.1 | 26973793 | 27100494 |
| *ANLN* | Non-kinase interactor | Proximal interactor | 7 | p14.2 | 36389821 | 36453791 |
| *ARFIP2* | Non-kinase interactor | Proximal interactor | 11 | p15.4 | 6474683 | 6481479 |
| *ARL13B* | Non-kinase interactor | Proximal interactor | 3 | q11.1 | 93980139 | 94055678 |
| *ARMCX3* | Non-kinase interactor | Proximal interactor | X | q22.1 | 101622797 | 101627843 |
| *ARPC1A* | Non-kinase interactor | Proximal interactor | 7 | q22.1 | 99325898 | 99366262 |
| *ARPC1B* | Non-kinase interactor | Proximal interactor | 7 | q22.1 | 99374249 | 99394816 |
| *ARPC2* | Non-kinase interactor | Proximal interactor | 2 | q35 | 218217141 | 218254356 |
| *ARPC3* | Non-kinase interactor | Proximal interactor | 12 | q24.11 | 110434823 | 110450422 |
| *ARPC4* | Non-kinase interactor | Proximal interactor | 3 | p25.3 | 9792495 | 9807101 |
| *ARPC5* | Non-kinase interactor | Proximal interactor | 1 | q25.3 | 183620846 | 183635783 |
| *ARPC5L* | Non-kinase interactor | Proximal interactor | 9 | q33.3 | 124862130 | 124877733 |
| *ATP6AP1* | Non-kinase interactor | Proximal interactor | X | q28 | 154428633 | 154436516 |
| *BAIAP2* | Non-kinase interactor | Proximal interactor | 17 | q25.3 | 81035122 | 81117434 |
| *BAIAP2L1* | Non-kinase interactor | Proximal interactor | 7 | q22.1 | 98291650 | 98401090 |
| *BASP1* | Non-kinase interactor | Proximal interactor | 5 | p15.1 | 17065598 | 17276843 |
| *BCAP31* | Non-kinase interactor | Proximal interactor | X | q28 | 153700492 | 153724565 |
| *BRK1* | Non-kinase interactor | Proximal interactor | 3 | p25.3 | 10115675 | 10127190 |

| Symbol | Subgroup | Type of protein | Chr | Band | Start | End |
|---|---|---|---|---|---|---|
| *C1QBP* | Non-kinase interactor | Proximal interactor | 17 | p13.2 | 5432777 | 5448830 |
| *CAPZB* | Non-kinase interactor | Proximal interactor | 1 | p36.13 | 19338775 | 19485539 |
| *CAV1* | Non-kinase interactor | Proximal interactor | 7 | q31.2 | 116524994 | 116561179 |
| *CAVIN1* | Non-kinase interactor | Proximal interactor | 17 | q21.2 | 42402449 | 42423256 |
| *CCDC115* | Non-kinase interactor | Proximal interactor | 2 | q21.1 | 130337933 | 130342699 |
| *CCDC187* | Non-kinase interactor | Proximal interactor | 9 | q34.3 | 136249973 | 136306901 |
| *CCDC88A* | Non-kinase interactor | Proximal interactor | 2 | p16.1 | 55287842 | 55419895 |
| *CCP110* | Non-kinase interactor | Proximal interactor | 16 | p12.3 | 19523811 | 19553408 |
| *CCT2* | Non-kinase interactor | Proximal interactor | 12 | q15 | 69585426 | 69601570 |
| *CCT6A* | Non-kinase interactor | Proximal interactor | 7 | p11.2 | 56051685 | 56063989 |
| *CCT7* | Non-kinase interactor | Proximal interactor | 2 | p13.2 | 73233420 | 73253021 |
| *CDC42EP1* | Non-kinase interactor | Proximal interactor | 22 | q13.1 | 37560480 | 37569405 |
| *CDC42EP2* | Non-kinase interactor | Proximal interactor | 11 | q13.1 | 65314866 | 65322417 |
| *CDC42EP3* | Non-kinase interactor | Proximal interactor | 2 | p22.2 | 37641882 | 37738468 |
| *CDC42EP4* | Non-kinase interactor | Proximal interactor | 17 | q25.1 | 73283624 | 73312005 |
| *CDC42EP5* | Non-kinase interactor | Proximal interactor | 19 | q13.42 | 54465026 | 54473296 |
| *CDC42SE1* | Non-kinase interactor | Proximal interactor | 1 | q21.3 | 151050971 | 151070325 |
| *CDC42SE2* | Non-kinase interactor | Proximal interactor | 5 | q31.1 | 131245493 | 131398447 |
| *CEP97* | Non-kinase interactor | Proximal interactor | 3 | q12.3 | 101724593 | 101770562 |
| *CKAP4* | Non-kinase interactor | Proximal interactor | 12 | q23.3 | 106237881 | 106304279 |
| *CLTC* | Non-kinase interactor | Proximal interactor | 17 | q23.1 | 59619689 | 59696956 |
| *COPS2* | Non-kinase interactor | Proximal interactor | 15 | q21.1 | 49106068 | 49155661 |
| *COPS4* | Non-kinase interactor | Proximal interactor | 4 | q21.22 | 83034447 | 83075818 |
| *CPD* | Non-kinase interactor | Proximal interactor | 17 | q11.2 | 30378927 | 30469989 |
| *CPNE2* | Non-kinase interactor | Proximal interactor | 16 | q13 | 57092583 | 57148369 |
| *CPSF7* | Non-kinase interactor | Proximal interactor | 11 | q12.2 | 61402641 | 61430031 |

| Symbol | Subgroup | Type of protein | Chr | Band | Start | End |
|---|---|---|---|---|---|---|
| *CYFIP1* | Non-kinase interactor | Proximal interactor | 15 | q11.2 | 22867052 | 22981063 |
| *DBN1* | Non-kinase interactor | Proximal interactor | 5 | q35.3 | 177456608 | 177474401 |
| *DBT* | Non-kinase interactor | Proximal interactor | 1 | p21.2 | 100186919 | 100249834 |
| *DDRGK1* | Non-kinase interactor | Proximal interactor | 20 | p13 | 3190350 | 3204685 |
| *DDX39B* | Non-kinase interactor | Proximal interactor | 6 | p21.33 | 31530219 | 31542448 |
| *DDX4* | Non-kinase interactor | Proximal interactor | 5 | q11.2 | 55738017 | 55817157 |
| *DIAPH1* | Non-kinase interactor | Direct effector | 5 | q31.3 | 141515016 | 141619055 |
| *DIAPH2* | Non-kinase interactor | Direct effector | X | q21.33 | 96684663 | 97604997 |
| *DIAPH3* | Non-kinase interactor | Direct effector | 13 | q21.2 | 59665583 | 60163928 |
| *DLG5* | Non-kinase interactor | Proximal interactor | 10 | q22.3 | 77790791 | 77926755 |
| *DSG1* | Non-kinase interactor | Proximal interactor | 18 | q12.1 | 31318160 | 31359246 |
| *DSG2* | Non-kinase interactor | Proximal interactor | 18 | q12.1 | 31498177 | 31549008 |
| *DSP* | Non-kinase interactor | Proximal interactor | 6 | p24.3 | 7541617 | 7586714 |
| *DST* | Non-kinase interactor | Proximal interactor | 6 | p12.1 | 56457987 | 56954830 |
| *EFHD2* | Non-kinase interactor | Proximal interactor | 1 | p36.21 | 15409888 | 15430339 |
| *ELMO2* | Non-kinase interactor | Proximal interactor | 20 | q13.12 | 46366050 | 46432985 |
| *EMC3* | Non-kinase interactor | Proximal interactor | 3 | p25.3 | 9962537 | 10011116 |
| *EMD* | Non-kinase interactor | Proximal interactor | X | q28 | 154379273 | 154381574 |
| *EPSTI1* | Non-kinase interactor | Proximal interactor | 13 | q14.11 | 42886388 | 42992271 |
| *ERBIN* | Non-kinase interactor | Proximal interactor | 5 | q12.3 | 65926556 | 66082546 |
| *ESYT1* | Non-kinase interactor | Proximal interactor | 12 | q13.2 | 56118250 | 56144674 |
| *FAF2* | Non-kinase interactor | Proximal interactor | 5 | q35.2 | 176447628 | 176510074 |
| *FAM65A* | Non-kinase interactor | Proximal interactor | 16 | q22.1 | 67552321 | 67580961 |
| *FAM65B* | Non-kinase interactor | Proximal interactor | 6 | p11.1 | 24797601 | 25042238 |

| Symbol | Subgroup | Type of protein | Chr | Band | Start | End |
|---|---|---|---|---|---|---|
| *FAM65C* | Non-kinase interactor | Proximal interactor | 20 | q13.13 | 49202645 | 49308065 |
| *FAM83B* | Non-kinase interactor | Proximal interactor | 6 | p12.1 | 54846771 | 54945099 |
| *FAM91A1* | Non-kinase interactor | Proximal interactor | 8 | q24.13 | 123768439 | 123815452 |
| *FAM135A* | Non-kinase interactor | Proximal interactor | 6 | q13 | 70412941 | 70561174 |
| *FAM169A* | Non-kinase interactor | Proximal interactor | 5 | q13.3 | 74777574 | 74866966 |
| *FERMT2* | Non-kinase interactor | Proximal interactor | 14 | q22.1 | 52857268 | 52952435 |
| *FLOT1* | Non-kinase interactor | Proximal interactor | 6 | p21.33 | 30727709 | 30742732 |
| *FLOT2* | Non-kinase interactor | Proximal interactor | 17 | q11.2 | 28879335 | 28897733 |
| *FMNL3* | Non-kinase interactor | Proximal interactor | 12 | q13.12 | 49636499 | 49708165 |
| *FNBP1* | Non-kinase interactor | Proximal interactor | 9 | q34.11 | 129887187 | 130043194 |
| *FNBP1L* | Non-kinase interactor | Proximal interactor | 1 | p22.1 | 93448118 | 93554661 |
| *GARRE1* | Non-kinase interactor | Proximal interactor | 19 | q13.11 | 34745442 | 34846491 |
| *GFOD1* | Non-kinase interactor | Proximal interactor | 6 | p23 | 13357830 | 13487662 |
| *GIT1* | Non-kinse interactor | Proximal interactor | 17 | q11.2 | 29573475 | 29594054 |
| *GIT2* | Non-kinse interactor | Proximal interactor | 12 | q24.11 | 109929802 | 109996389 |
| *GJA1* | Non-kinase interactor | Proximal interactor | 6 | q22.31 | 121435595 | 121449727 |
| *GOLGA3* | Non-kinase interactor | Proximal interactor | 12 | q24.33 | 132768914 | 132829078 |
| *GOLGA8R* | Non-kinase interactor | Proximal interactor | 15 | q13.2 | 30403740 | 30414162 |
| *GPS1* | Non-kinase interactor | Proximal interactor | 17 | q25.3 | 82050691 | 82057470 |
| *HGS* | Non-kinase interactor | Proximal interactor | 17 | q25.3 | 81683326 | 81703138 |
| *HINT2* | Non-kinase interactor | Proximal interactor | 9 | p13.3 | 35812960 | 35815354 |
| *HMOX2* | Non-kinase interactor | Proximal interactor | 16 | p13.3 | 4474690 | 4510347 |
| *HNRNPC* | Non-kinase interactor | Proximal interactor | 14 | q11.2 | 21209136 | 21269494 |
| *HSPE1* | Non-kinase interactor | Proximal interactor | 2 | q33.1 | 197500140 | 197503449 |
| *IQGAP1* | Non-kinase interactor | Proximal interactor | 15 | q26.1 | 90388242 | 90502239 |
| *IQGAP2* | Non-kinase interactor | Proximal interactor | 5 | q13.3 | 76403285 | 76708132 |

| Symbol | Subgroup | Type of protein | Chr | Band | Start | End |
|---|---|---|---|---|---|---|
| *IQGAP3* | Non-kinase interactor | Proximal interactor | 1 | q22 | 156525405 | 156572604 |
| *ITGB1* | Non-kinase interactor | Proximal interactor | 10 | p11.22 | 32887273 | 33005792 |
| *JMY* | Non-kinase interactor | Proximal interactor | 5 | q14.1 | 79236131 | 79327211 |
| *JUP* | Non-kinase interactor | Proximal interactor | 17 | q21.2 | 41754604 | 41786931 |
| *KCTD3* | Non-kinase interactor | Proximal interactor | 1 | q41 | 215567304 | 215621807 |
| *KIDINS220* | Non-kinase interactor | Proximal interactor | 2 | p25.1 | 8721081 | 8837630 |
| *KIF14* | Non-kinase interactor | Proximal interactor | 1 | q32.1 | 200551497 | 200620751 |
| *KTN1* | Non-kinase interactor | Proximal interactor | 14 | q22.3 | 55559072 | 55701526 |
| *LAMTOR1* | Non-kinase interactor | Proximal interactor | 11 | q13.4 | 72085895 | 72103297 |
| *LBR* | Non-kinase interactor | Proximal interactor | 1 | q42.12 | 225401502 | 225428925 |
| *LEMD3* | Non-kinase interactor | Proximal interactor | 12 | q14.3 | 65169583 | 65248355 |
| *LETM1* | Non-kinase interactor | Proximal interactor | 4 | p16.3 | 1811479 | 1856156 |
| *LMAN1* | Non-kinase interactor | Proximal interactor | 18 | q21.32 | 59327823 | 59359265 |
| *LMNB1* | Non-kinase interactor | Proximal interactor | 5 | q23.2 | 126776623 | 126837020 |
| *LRRC1* | Non-kinase interactor | Proximal interactor | 6 | p12.1 | 53794497 | 53924125 |
| *MACO1* | Non-kinase interactor | Proximal interactor | 1 | p36.11 | 25430858 | 25500209 |
| *MCAM* | Non-kinase interactor | Proximal interactor | 11 | q23.3 | 119308529 | 119321521 |
| *MOSPD2* | Non-kinase interactor | Proximal interactor | X | p22.2 | 14873421 | 14922327 |
| *MPP7* | Non-kinase interactor | Proximal interactor | 10 | p12.1 | 28050993 | 28334486 |
| *MPRIP* | Non-kinase interactor | Proximal interactor | 17 | p11.2 | 17042457 | 17217679 |
| *MTMR1* | Non-kinase interactor | Proximal interactor | X | q28 | 150692971 | 150765108 |
| *MTR* | Non-kinase interactor | Proximal interactor | 1 | q43 | 236795260 | 236921278 |
| *MUC13* | Non-kinase interactor | Proximal interactor | 3 | q21.2 | 124905442 | 124953819 |
| *MYH11* | Non-kinase interactor | Proximal interactor | 16 | p13.11 | 15703135 | 15857028 |
| *MYL12B* | Non-kinase interactor | Proximal interactor | 18 | p11.31 | 3261479 | 3278461 |
| *MYO6* | Non-kinase interactor | Proximal interactor | 6 | q14.1 | 75749192 | 75919537 |

| Symbol | Subgroup | Type of protein | Chr | Band | Start | End |
|---|---|---|---|---|---|---|
| *NAP1L1* | Non-kinase interactor | Proximal interactor | 12 | q21.2 | 76036585 | 76084735 |
| *NCF2* | Non-kinase interactor | Proximal interactor | 1 | q25.3 | 183555562 | 183590876 |
| *NCK1* | Non-kinase interactor | Proximal interactor | 3 | q22.3 | 136862208 | 136951606 |
| *NCK2* | Non-kinase interactor | Proximal interactor | 2 | q12.2 | 105744912 | 105894274 |
| *NCKAP1* | Non-kinase interactor | Proximal interactor | 2 | q32.1 | 182909115 | 183038858 |
| *NDUFA5* | Non-kinase interactor | Proximal interactor | 7 | q31.32 | 123536997 | 123557904 |
| *NDUFS3* | Non-kinase interactor | Proximal interactor | 11 | p11.2 | 47565336 | 47584562 |
| *NHS* | Non-kinase interactor | Proximal interactor | X | p22.2 | 17375200 | 17735994 |
| *NIPSNAP2* | Non-kinase interactor | Proximal interactor | 7 | p11.2 | 55951793 | 56000181 |
| *NISCH* | Non-kinase interactor | Proximal interactor | 3 | p21.1 | 52455118 | 52493068 |
| *NOC2L* | Non-kinase interactor | Proximal interactor | 1 | p36.33 | 944203 | 959309 |
| *NSFL1C* | Non-kinase interactor | Proximal interactor | 20 | p13 | 1442162 | 1473842 |
| *NUDC* | Non-kinase interactor | Proximal interactor | 1 | p36.11 | 26900238 | 26946871 |
| *OSBPL11* | Non-kinase interactor | Proximal interactor | 3 | q21.2 | 125528858 | 125595497 |
| *PAK1IP1* | Non-kinase interactor | Proximal interactor | 6 | p24.2 | 10694972 | 10709782 |
| *PCDH7* | Non-kinase interactor | Proximal interactor | 4 | p15.1 | 30720415 | 31146805 |
| *PEAK1* | Non-kinase interactor | Proximal interactor | 15 | q24.3 | 77100656 | 77420144 |
| *PGRMC2* | Non-kinase interactor | Proximal interactor | 4 | q28.2 | 128269237 | 128288829 |
| *PHIP* | Non-kinase interactor | Proximal interactor | 6 | q14.1 | 78934419 | 79078254 |
| *PKP4* | Non-kinase interactor | Proximal interactor | 2 | q24.1 | 158456952 | 158682879 |
| *PLD1* | Non-kinase interactor | Proximal interactor | 3 | q26.31 | 171600404 | 171810950 |
| *PLD2* | Non-kinase interactor | Proximal interactor | 17 | p13.2 | 4807152 | 4823434 |
| *POTEE* | Non-kinase interactor | Proximal interactor | 2 | q21.1 | 131209536 | 131265278 |
| *PTPN13* | Non-kinase interactor | Proximal interactor | 4 | q21.3 | 86594315 | 86815171 |
| *RAB7A* | Non-kinase interactor | Proximal interactor | 3 | q21.3 | 128693669 | 128825942 |
| *RASAL2* | Non-kinase interactor | Proximal interactor | 1 | q25.2 | 178094104 | 178484147 |

| Symbol | Subgroup | Type of protein | Chr | Band | Start | End |
|---|---|---|---|---|---|---|
| *RBBP6* | Non-kinase interactor | Proximal interactor | 16 | p12.1 | 24537693 | 24572863 |
| *RBMX* | Non-kinase interactor | Proximal interactor | X | q26.3 | 136848004 | 136880764 |
| *RHPN1* | Non-kinase interactor | Proximal interactor | 8 | q24.3 | 143368876 | 143384221 |
| *RHPN2* | Non-kinase interactor | Proximal interactor | 19 | q13.11 | 32978592 | 33064888 |
| *RNF20* | Non-kinase interactor | Proximal interactor | 9 | q31.1 | 101533853 | 101563344 |
| *RRAS2* | Non-kinase interactor | Proximal interactor | 11 | p15.2 | 14277922 | 14364506 |
| *RTKN* | Non-kinase interactor | Proximal interactor | 2 | p13.1 | 74425835 | 74442422 |
| *RTKN2* | Non-kinase interactor | Proximal interactor | 10 | q21.2 | 62183035 | 62268844 |
| *SCFD1* | Non-kinase interactor | Proximal interactor | 14 | q12 | 30622291 | 30737694 |
| *SCRIB* | Non-kinase interactor | Proximal interactor | 8 | q24.3 | 143790920 | 143815773 |
| *SEMA4F* | Non-kinase interactor | Proximal interactor | 2 | p13.1 | 74654228 | 74683853 |
| *SENP1* | Non-kinase interactor | Proximal interactor | 12 | q13.11 | 48042897 | 48106079 |
| *SH3PXD2A* | Non-kinase interactor | Proximal interactor | 10 | q24.33 | 103594027 | 103855543 |
| *SH3RF1* | Non-kinase interactor | Proximal interactor | 4 | q33 | 169094259 | 169270956 |
| *SH3RF2* | Non-kinase interactor | Proximal interactor | 5 | q32 | 145936578 | 146081791 |
| *SH3RF3* | Non-kinase interactor | Proximal interactor | 2 | q13 | 109129205 | 109504634 |
| *SHKBP1* | Non-kinase interactor | Proximal interactor | 19 | q13.2 | 40576853 | 40591399 |
| *SHMT2* | Non-kinase interactor | Proximal interactor | 12 | q13.3 | 57229573 | 57234935 |
| *SLC1A5* | Non-kinase interactor | Proximal interactor | 19 | q13.32 | 46774883 | 46788594 |
| *SLC4A7* | Non-kinase interactor | Proximal interactor | 3 | p24.1 | 27372721 | 27484420 |
| *SLITRK3* | Non-kinase interactor | Proximal interactor | 3 | q26.1 | 165186720 | 165197109 |
| *SLITRK5* | Non-kinase interactor | Proximal interactor | 13 | q31.2 | 87671371 | 87696272 |
| *SNAP23* | Non-kinase interactor | Proximal interactor | 15 | q15.1 | 42491233 | 42545356 |
| *SOWAHC* | Non-kinase interactor | Proximal interactor | 2 | q13 | 109614364 | 109618990 |
| *SOX9* | Non-kinase interactor | Proximal interactor | 17 | q24.3 | 72121020 | 72126416 |
| *SPEN* | Non-kinase interactor | Proximal interactor | 1 | p36.21 | 15836095 | 15940456 |

| Symbol | Subgroup | Type of protein | Chr | Band | Start | End |
|---|---|---|---|---|---|---|
| *SPTAN1* | Non-kinase interactor | Proximal interactor | 9 | q34.11 | 128552558 | 128633662 |
| *SPTBN1* | Non-kinase interactor | Proximal interactor | 2 | p16.2 | 54456317 | 54671446 |
| *SRA1* | Non-kinase interactor | proximal interactor | 5 | q31.3 | 140537340 | 140557677 |
| *SRRM1* | Non-kinase interactor | proximal interactor | 1 | p36.11 | 24631716 | 24673281 |
| *STAM* | Non-kinase interactor | proximal interactor | 10 | p12.33 | 17644151 | 17716824 |
| *STAM2* | Non-kinase interactor | proximal interactor | 2 | q23.3 | 152116801 | 152175763 |
| *STBD1* | Non-kinase interactor | proximal interactor | 4 | q21.1 | 76306733 | 76311130 |
| *STEAP3* | Non-kinase interactor | proximal interactor | 2 | q14.2 | 119223831 | 119265652 |
| *STOM* | Non-kinase interactor | proximal interactor | 9 | q33.2 | 121338987 | 121370304 |
| *STX5* | Non-kinase interactor | proximal interactor | 11 | q12.3 | 62806860 | 62832051 |
| *SWAP70* | Non-kinase interactor | proximal interactor | 11 | p15.4 | 9664077 | 9752993 |
| *TEX2* | Non-kinase interactor | proximal interactor | 17 | q23.3 | 64147227 | 64263260 |
| *TFRC* | Non-kinase interactor | proximal interactor | 3 | q29 | 196027183 | 196082096 |
| *TJP2* | Non-kinase interactor | proximal interactor | 9 | q21.11 | 69121264 | 69274615 |
| *TMEM59* | Non-kinase interactor | proximal interactor | 1 | p32.3 | 54026681 | 54053504 |
| *TMEM87A* | Non-kinase interactor | proximal interactor | 15 | q15.1 | 42210447 | 42273584 |
| *TMOD3* | Non-kinase interactor | proximal interactor | 15 | q21.2 | 51829628 | 51947295 |
| *TMPO* | Non-kinase interactor | proximal interactor | 12 | q23.1 | 98515579 | 98550351 |
| *TOR1AIP1* | Non-kinase interactor | proximal interactor | 1 | q25.2 | 179882042 | 179920077 |
| *TPM3* | Non-kinase interactor | proximal interactor | 1 | q21.3 | 154155304 | 154194648 |
| *TPM4* | Non-kinase interactor | proximal interactor | 19 | p13.12 | 16067021 | 16103002 |
| *TRA2B* | Non-kinase interactor | proximal interactor | 3 | q27.2 | 185914558 | 185938103 |
| *TRIOBP* | Non-kinase interactor | proximal interactor | 22 | q13.1 | 37697048 | 37776556 |
| *TRIP10* | Non-kinase interactor | proximal interactor | 19 | p13.3 | 6737925 | 6751530 |
| *TUBA1B* | Non-kinase interactor | proximal interactor | 12 | q13.12 | 49127782 | 49131397 |
| *TWF1* | Non-kinase interactor | proximal interactor | 12 | q12 | 43793723 | 43806328 |

| Symbol | Subgroup | Type of protein | Chr | Band | Start | End |
|---|---|---|---|---|---|---|
| *TXNL1* | Non-kinase interactor | proximal interactor | 18 | q21.31 | 56597209 | 56651600 |
| *UACA* | Non-kinase interactor | proximal interactor | 15 | q23 | 70654554 | 70763558 |
| *UHRF1BP1L* | Non-kinase interactor | Proximal interactor | 12 | q23.1 | 100028455 | 100142874 |
| *USP9X* | Non-kinase interactor | Proximal interactor | X | p11.4 | 41085445 | 41236579 |
| *VAMP3* | Non-kinase interactor | Proximal interactor | 1 | p36.23 | 7771296 | 7781432 |
| *VANGL1* | Non-kinase interactor | Proximal interactor | 1 | p13.1 | 115641970 | 115698224 |
| *VANGL2* | Non-kinase interactor | Proximal interactor | 1 | q23.2 | 160400564 | 160428670 |
| *VAPB* | Non-kinase interactor | Proximal interactor | 20 | q13.32 | 58389229 | 58451101 |
| *VCP* | Non-kinase interactor | Proximal interactor | 9 | p13.3 | 35053928 | 35072668 |
| *VIM* | Non-kinase interactor | Proximal interactor | 10 | p13 | 17228241 | 17237593 |
| *WAS* | Non-kinase interactor | Proximal interactor | X | p11.23 | 48676596 | 48691427 |
| *WASF1* | Non-kinase interactor | Proximal interactor | 6 | q21 | 110099819 | 110180004 |
| *WASF2* | Non-kinase interactor | Proximal interactor | 1 | p36.11 | 27404230 | 27490167 |
| *WASF3* | Non-kinase interactor | Proximal interactor | 13 | q12.13 | 26557683 | 26688948 |
| *WASL* | Non-kinase interactor | Proximal interactor | 7 | q31.32 | 123681943 | 123749003 |
| *WDR11* | Non-kinase interactor | Proximal interactor | 10 | q26.12 | 120851305 | 120909524 |
| *WDR6* | Non-kinase interactor | Proximal interactor | 3 | p21.31 | 49007062 | 49015953 |
| *WDR81* | Non-kinase interactor | Proximal interactor | 17 | p13.3 | 1716523 | 1738599 |
| *WDR91* | Non-kinase interactor | Proximal interactor | 7 | q33 | 135183839 | 135211555 |
| *WHAMM* | Non-kinase interactor | Proximal interactor | 15 | q25.2 | 82809628 | 82836108 |
| *WIPF1* | Non-kinase interactor | Proximal interactor | 2 | q31.1 | 174559572 | 174682916 |
| *WIPF2* | Non-kinase interactor | Proximal interactor | 17 | q21.2 | 40219304 | 40284136 |
| *WIPF3* | Non-kinase interactor | Proximal interactor | 7 | p14.3 | 29806486 | 29917066 |
| *WWP2* | Non-kinase interactor | Proximal interactor | 16 | q22.1 | 69762328 | 69941741 |
| *YKT6* | Non-kinase interactor | Proximal interactor | 7 | p13 | 44200959 | 44214294 |
| *ZNF512B* | Non-kinase interactor | Proximal interactor | 20 | q13.33 | 63956704 | 63969930 |

* **Direct effector,** downstream element that can physically interact with GTPases; **distal effector,** RHO downstream signaling element that is located further downstream of the proximal effectors; **proximal interactor,** protein belonging to the large-scale interactome of specific RHO GTPases according to proteomics determinations.

### TOTAL NUMBER OF GENES/PROTEINS

| Type | | Number | Total |
|---|---|---|---|
| RHO GTPases | | 23 (including 3 RHOBTBs) | 20 |
| RHO GAP domain containing proteins | | 70 (including 2 with DH domains) | 69 |
| RHO GEFs or contaning similar structural domains | DH family members | 75 (including 2 with GAP domains). Total 61 genes | 87 (85 genes) |
| | DOCK family members | 11 | |
| | Armadillo domain | 1 | |
| RHO GDI | | 3 | 3 |
| Downstream elements or interactors with kinase activity | | 64 | 64 |
| Downstream elements or interactors lacking kinase activity | | 243 | 243 |
| **Total** | | | **484** |
