## Supplementary Table 2 for "PAN-CANCER ANALYSES IDENTIFY ONCOGENIC DRIVERS, EXPRESSION SIGNATURES, AND THERAPEUTIC VULNERABILITIES IN RHO GTPase PATHWAY GENES"

**Supplementary Table 2. Abbreviations for cancer subtypes used in this study**

| Abbreviation | Cancer type |
|---|---|
| LAML | Acute myeloid leukemia |
| ACC | Adrenocortical carcinoma |
| BLCA | Bladder urothelial carcinoma |
| LGG | Brain lower grade glioma |
| BRCA | Breast invasive carcinoma |
| CESC | Cervical squamous cell carcinoma and endocervical adenocarcinoma |
| CHOL | Cholangiocarcinoma |
| COAD | Colon adenocarcinoma |
| ESCA | Esophageal carcinoma |
| GBM | Glioblastoma multiforme |
| HNSC | Head and neck squamous cell carcinoma |
| KICH | Kidney chromophobe |
| KIRC | Kidney renal clear cell carcinoma |
| KIRP | Kidney renal papillary cell carcinoma |
| LIHC | Liver hepatocellular carcinoma |
| LUAD | Lung adenocarcinoma |
| LUSC | Lung squamous cell carcinoma |
| DLBC | Lymphoid neoplasm diffuse large B-cell lymphoma |
| MESO | Mesothelioma |
| OV | Ovarian serous cystadenocarcinoma |
| PAAD | Pancreatic adenocarcinoma |
| PCPG | Pheochromocytoma and paraganglioma |
| PRAD | Prostate adenocarcinoma |
| READ | Rectum adenocarcinoma |
| SARC | Sarcoma |
| SKCM | Skin cutaneous melanoma |
| STAD | Stomach adenocarcinoma |
| TGCT | Testicular germ cell tumors |
| THYM | Thymoma |
| THCA | Thyroid carcinoma |
| UCS | Uterine carcinosarcoma |
| UCEC | Uterine corpus endometrial carcinoma |
| UVM | Uveal melanoma |
