## Supplementary Table 3 for "PAN-CANCER ANALYSES IDENTIFY ONCOGENIC DRIVERS, EXPRESSION SIGNATURES, AND THERAPEUTIC VULNERABILITIES IN RHO GTPase PATHWAY GENES"

### Supplementary Table S3. List of RHO GTPase pathway genes with alterations found in cancer

| Symbol | Subgroup* | Change** |
|---|---|---|
| *RAC1**** | RHO GTPase | PS, HS, es |
| *RHOA* | RHO GTPase | PS, HS, es |
| *RHOB* | RHO GTPase | PS, HS |
| *RND1* | RHO GTPase | HS |

| Symbol | Subgroup | Change |
|---|---|---|
| *ARAP2* | RHO GAP domain | HS |
| *ARHGAP1* | RHO GAP domain | HS |
| *ARHGAP9* | RHO GAP domain | HS |
| *ARHGAP29* | RHO GAP domain | HS |
| *ARHGAP30* | RHO GAP domain | HS |
| *ARHGAP35* | RHO GAP domain | PS, HS |
| *ARHGAP39* | RHO GAP domain | Amp |

| Symbol | Subgroup | Change |
|---|---|---|
| *CHN2* | RHO GAP domain | HS |
| *DEPDC1* | RHO GAP domain | Amp |
| *DEPDC1B* | RHO GAP domain | HS |
| *PIK3R1* | RHO GAP domain | PS, HS |
| *PIK3R2* | RHO GAP domain | HS |
| *STARD13* | RHO GAP domain | HS |
| *ARHGEF15* | RHO GEF (DH) | HS |
| *ARHGEF25* | RHO GEF (DH) | HS |
| *ECT2* | RHO GEF (DH) | Amp, ES, es |
| *FGD5* | RHO GEF (DH) | HS |
| *MCF2* | RHO GEF (DH) | HS |
| *PLEKHG2* | RHO GEF (DH) | HS |
| *PLEKHG4* | RHO GEF (DH) | HS |

| Symbol | Subgroup | Change |
|---|---|---|
| *PLEKHG4B* | RHO GEF (DH) | Amp |
| *PLEKHG7* | RHO GEF (DH) | HS |
| *DOCK1* | RHO GEF (DOCK) | HS |

| Symbol | Subgroup | Type of protein | Change |
|---|---|---|---|
| *CDC42BPA* | Kinase interactor | Direct effector* | HS |
| *MAP3K1* | Kinase interactor | Direct effector | PS |
| *MYLK* | Kinase interactor | Direct effector | HS |
| *PAK6* | Kinase interactor | Direct effector | HS |
| *PIK3CA* | Kinase interactor | Direct effector | PS, HS |
| *PKN2* | Kinase interactor | Direct effector | HS |
| *PLK1* | Kinase interactor | Distal effector | HS, UP, ES |
| *RPS6KB1* | Kinase interactor | Distal effector | HS |
| *STK10* | Kinase interactor | Distal effector | HS |
| *TNK1* | Kinase interactor | Direct effector | HS |
| *ACTB* | Non-kinase interactor | Direct effector | PS, HS |
| *ACTC1* | Non-kinase interactor | Direct effector | HS |

| Symbol | Subgroup | Type of protein | Change |
|---|---|---|---|
| *ACTR3B* | Non-kinase interactor | Direct effector | HS |
| *ANLN* | Non-kinase interactor | Proximal interactor | Amp, ES |
| *ARL13B* | Non-kinase interactor | Proximal interactor | HS |
| *ARPC1A* | Non-kinase interactor | Proximal interactor | HS |
| *CKAP4* | Non-kinase interactor | Proximal interactor | HS |
| *DST* | Non-kinase interactor | Proximal interactor | HS |

| Symbol | Subgroup | Type of protein | Change |
|---|---|---|---|
| *FAM65C* | Non-kinase interactor | Proximal interactor | HS |
| *GFOD1* | Non-kinase interactor | Proximal interactor | HS |
| *IQGAP3* | Non-kinase interactor | Proximal interactor | HS |
| *ITGB1* | Non-kinase interactor | Proximal interactor | HS |
| *KIF14* | Non-kinase interactor | Proximal interactor | Amp, ES |
| *MTMR1* | Non-kinase interactor | Proximal interactor | HS |
| *MYH11* | Non-kinase interactor | Proximal interactor | HS, Del |
| *NHS* | Non-kinase interactor | Proximal interactor | HS |
| *OSBPL11* | Non-kinase interactor | Proximal interactor | HS |
| *PAK1IP1* | Non-kinase interactor | Proximal interactor | HS |

| Symbol | Subgroup | Type of protein | Change |
|---|---|---|---|
| *POTEE* | Non-kinase interactor | Proximal interactor | HS |
| *PTPN13* | Non-kinase interactor | Proximal interactor | HS |
| *RHPN1* | Non-kinase interactor | Proximal interactor | Amp |
| *RRAS2* | Non-kinase interactor | Proximal interactor | HS |
| *SH3RF2* | Non-kinase interactor | Proximal interactor | HS |
| *SLC4A7* | Non-kinase interactor | Proximal interactor | HS |
| *SLITRK3* | Non-kinase interactor | Proximal interactor | HS |
| *SOX9* | Non-kinase interactor | Proximal interactor | PS, ES |
| *SPTAN1* | Non-kinase interactor | Proximal interactor | PS |
| *TFRC* | Non-kinase interactor | proximal interactor | Amp, es |
| *TRA2B* | Non-kinase interactor | proximal interactor | HS |
| *TXNL1* | Non-kinase interactor | proximal interactor | HS |
| *WASL* | Non-kinase interactor | Proximal interactor | HS |
| *WIPF1* | Non-kinase interactor | Proximal interactor | HS |

* **Direct effector,** downstream element that can physically interact with GTPases; **distal effector,** RHO downstream signaling element that is located further downstream of the proximal effectors; **proximal interactor,** protein belonging to the large-scale interactome of specific RHO GTPases according to proteomics determinations.

** **PS,** positively selected mutations; **HS,** hotspot mutations; **Amp,** amplified; **Del,** deleted; **ES,** important for proliferation in a large number of cancer cell lines; **es,** important for proliferation of specific cancer cell lines.
